# Emergent chirality in active solid rotation of pancreas spheres

**DOI:** 10.1101/2022.09.29.510101

**Authors:** Tzer Han Tan, Aboutaleb Amiri, Irene Seijo-Barandiarán, Michael F. Staddon, Anne Materne, Sandra Tomas, Charlie Duclut, Marko Popović, Anne Grapin-Botton, Frank Jülicher

**Affiliations:** Max Planck Institute for the Physics of Complex Systems, Dresden, Germany; Max Planck Institute of Molecular Cell Biology and Genetics, Dresden, Germany; Center for Systems Biology Dresden, Dresden, Germany; Cluster of Excellence Physics of Life, TU Dresden, Dresden, Germany; Center for Molecular and Cellular Bioengineering, TU Dresden, Dresden, Germany; Université Paris Cité, Laboratoire Matière et Systèmes Complexes (MSC), UMR 7057 CNRS, Paris, France

**Author notes:** Laboratoire Physico-Chimie Curie, CNRS UMR168, Institut Curie, Université PSL, Sorbonne Université, 75005, Paris, France. These authors contributed equally and are joint first authors (Tzer Han Tan, Aboutaleb Amiri, Irene Seijo-Barandiarán).

## Abstract

Collective cell dynamics play a crucial role in many developmental and physiological contexts. While two-dimensional (2D) cell migration has been widely studied, how three-dimensional (3D) geometry and topology interplay with collective cell behavior to determine dynamics and functions remains an open question. In this work, we elucidate the biophysical mechanism underlying rotation in spherical tissues, a phenomenon widely reported both *in vivo* and *in vitro*. Using murine pancreas-derived organoids as a model system, we find that epithelial spheres exhibit persistent rotation, rotational axis drift and rotation arrest. Using a 3D vertex model, we demonstrate how the interplay between traction force and polarity alignment can account for these distinct rotational dynamics. Furthermore, our analysis shows that the spherical tissue rotates as an active solid and exhibits spontaneous chiral symmetry breaking. Using a continuum model, we demonstrate how the types and location of topological defects in the polarity field underlie this symmetry breaking process. Altogether, our work shows that tissue chirality can arise via topological defects in the pattern of cell traction forces, with potential implications for left-right symmetry breaking processes in morphogenetic events.

## Main text

Collective cell migration is an important phenomenon in many biological systems [1], ranging from bacterial colonies [2, 3] to morphogenesis [4] and wound healing in multicellular organisms [5]. Through the interplay between directed motion, neighbor alignment and mechanical interactions, cell collectives exhibit emergent structures and dynamics that are crucial for their function. These include topological defects in 2D [6, 7] and 3D [8, 9] tissues during morphogenesis, jamming transition during vertebrate body axis elongation [10], and chiral tissue flows during gastrulation [11]. Recent work in synthetic active matter shows that curved surfaces and topological constraints can modulate the collective dynamics in active matter systems, resulting in shape changing vesicles [12], curvature-dependent defect unbinding [13] and topological sound [14]. This raises the question of how geometry and topology govern the emergence of collective patterns of cell migration from cellular properties and cell interactions.

A prominent phenomenon of collective cell migration on curved surfaces is tissue rotation in spherical geometry, which has been reported both *in vitro*, in the case of MDCK spheres [15, 16] and cancer spheroids [17], and *in vivo*, as in *Drosophila* egg chamber rotation [18]. Rotations were also recently described in organoid systems derived from primary cells from the mammary gland [19–22], indicating that tissue rotation is an intrinsic dynamics of many 3D multicellular structures. Yet, the biophysical mechanisms underlying tissue rotational dynamics and how different rotational modes can be biologically controlled remain open questions.

### Distinct rotational dynamics in pancreas spheres

Here, we use spherical pancreas organoids (henceforth ‘pancreas spheres’), derived from mouse pancreas progenitor cells [23] (see SI Sec. 1), as a model to investigate the collective cell motion in tissue rotation. These pancreas spheres are single-layer epithelia with apico-basal polarity. The apical side faces the fluid-filled lumen while the basal side faces outwards towards the Matrigel (which acts as extracellular matrix) (Fig. 1a). Live imaging experiments show that pancreas spheres derived from E13.5 mice exhibit a range of rotational dynamics. In the short term (of the order of hours), they either rotate (Fig. 1f(i), SI Video 1) or not rotate (Fig. 1f(ii), SI Video 2). At longer timescale (> 10 hours), more complex transitional dynamics are observed, such as rotation arrest (Fig. 1g(i), SI Video 3) and change in rotation axis (Fig. 1g(ii), SI Video 4).

**Figure 1:**
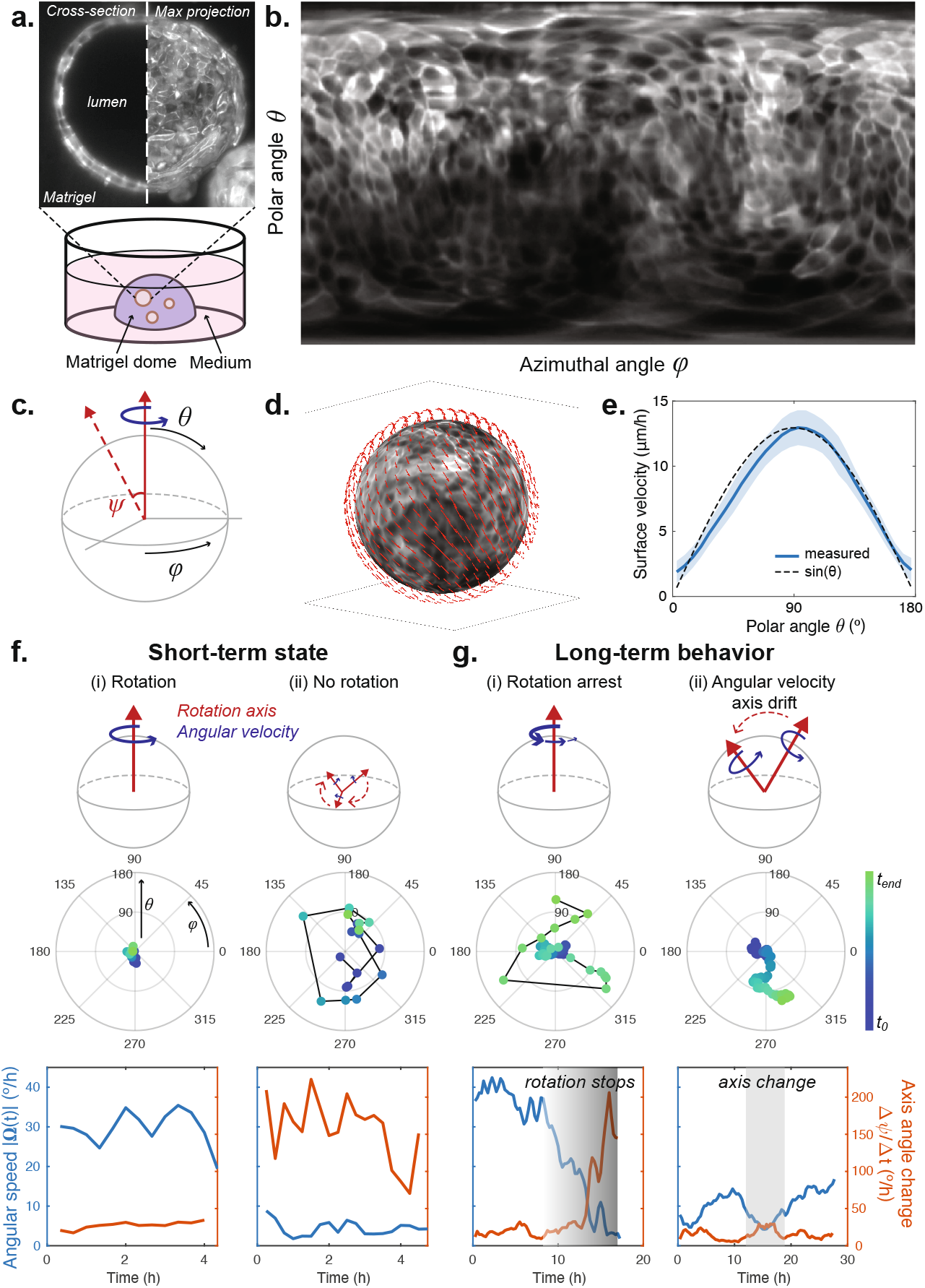
Distinct rotational dynamics in pancreas spheres. **a,** Experimental setup: pancreas progenitor cells form single layer epithelial spheres when cultured in Matrigel. **b,** Mercator projection of the apical surface of a pancreas sphere. **c,** Schematic defining the angular conventions: *ϕ* denotes azimuthal angle, *θ* the polar angle and *ψ* the angular change of rotation axis. **d,** Surface velocity of a rotating sphere as measured by particle image velocimetry (PIV). **e,** Surface velocity of a rotating sphere as a function of polar angle *θ*. **f-g,** Rotational dynamics that pancreas spheres exhibit in the short (**f**) and long (**g**) term. The first row shows a schematic for the type of rotational dynamics. The second row shows the angular position of the rotation axis as a function of time in the (*θ, ϕ*) polar plot. The third row shows the magnitude of the angular velocity and the rotational axis angular change. We show a representative case for each type of dynamics here. Additional data is included in SI Fig. 1.

To quantitatively characterize the complex rotational dynamics, we develop a computational pipeline to segment the spheres’ apical surface and visualize its fluorescence intensity by a Mercator projection (Fig. 1b, SI Sec. 2.2). By performing particle image velocimetry (PIV) analysis, we find that the resultant time-lapsed projection shows long range correlated tissue flow, with localized vortical motion that corresponds to the two poles of tissue rotation (SI Fig. 2, SI Video 5). This confirms that collective cell migration underlies the rotational motion of pancreas spheres.

To determine the angular velocity and the rotation axis, we reconstructed the full three-dimensional (3D) tissue flow field on the sphere (Fig. 1d) and computed its angular momentum (see SI Sec. 2.3). We find that the azimuthal velocity with respect to the instantaneous rotation axis follows a sine profile (Fig. 1e), suggesting that the pancreas spheres rotate akin to a solid sphere.

The dynamics of pancreas sphere observed on shorter time-scales up to 4 hours can be classified as either rotating (Fig. 1f(i)) or non-rotating (Fig. 1f(ii)). During persistent rotation, the angular speed remains roughly constant and the orientation of rotation axis remains stable. In contrast, when there is no rotation, the sphere is characterized by a small angular speed and large jumps in rotation axis orientation. We then imaged the dynamics of pancreas spheres on longer time scales up to 30 hours. This allowed us to capture transitions between the rotating and non-rotating states (Fig. 1g(i)). During a transition, the angular velocity slows down and the rotation axis becomes unstable, leading to a marked increase in axis angle change (Fig. 1g(i)). Furthermore, we have observed persistently rotating spheres that occasionally change their axis of rotation (Fig. 1g(ii)). This is characterized by a slow meandering in the orientation of rotation axis, coinciding with periods of lower angular velocity and higher rate of axis angle change.

### Solid and flowing regimes of active vertex model

These various forms of dynamics appear within the same culture condition, suggesting that an underlying biophysical mechanism is responsible for the observed types of dynamics and transitions between them. To identify this mechanism, we developed a mechanical model of pancreas spheres using a 3D vertex model. Vertex models have been used to study the mechanical aspects of epithelial morphogenesis that regulate their packing geometry [24], motility-driven rigidity transition [25], and developing tissues as amorphous solids that exhibit yielding transition under shear [26]. Due to the active nature of our system, we included two necessary ingredients: (1) traction force that allows the cells to move with respect to the ECM; and (2) cell polarity that determines the direction of the traction force.

Specifically, we extend the model in ref [24] to a spherical geometry and define the forces stemming from the mechanical properties of individual cells and the interaction between them based on the work function

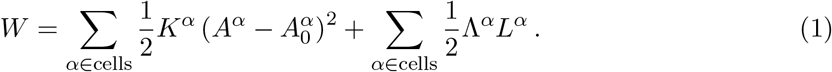

This model captures the cell shapes on the sphere surface as polygons. The first term accounts for cell area elasticity with 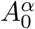 being the preferred cell area and *K^α^* the area stiffness. The second term accounts for cell bond tension, where Λ^*α*^ is the bond tension magnitude, and *L^α^* denotes cell perimeter [24, 27]. Here, we consider for simplicity the case where the mechanical parameters are uniform in the tissue and equal for all cells: 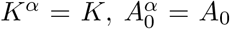 and Λ^*α*^ = Λ. At each vertex *m* located at position ***X**_m_*, force balance reads:

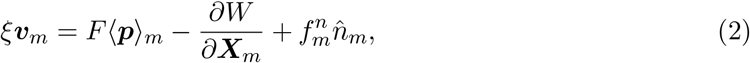

where ***v**_m_* is the velocity of vertex *m*, and *ξ* is the friction coefficient with the external environment. The first term on the right-hand side (r.h.s.) describes traction forces of magnitude *F* exerted by the (three) cells abutting at vertex *m* and in the average direction of their polarities 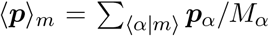, where 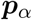 and *M_α_* are each cell’s polarity and number of vertices, re-spectively (see Fig. 2a). We consider a non-deforming spherical geometry by setting 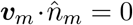 which specifies the normal force at vertex *m* with magnitude 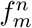 (see SI Sec. 3).

**Figure 2:**
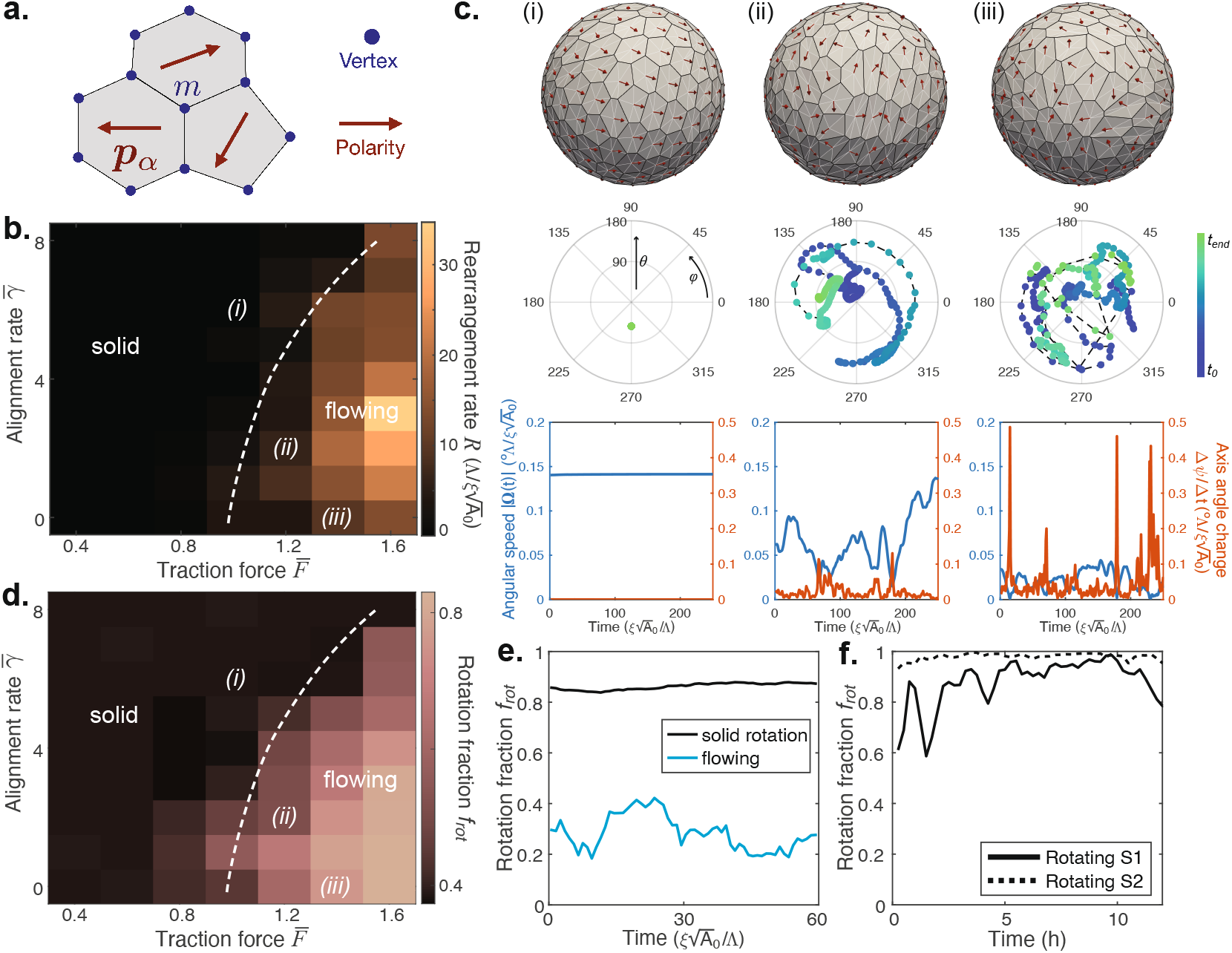
Solid and flowing regimes of the active vertex model. **a,** Schematic of the active vertex model. Cells are represented as polygons with vertices subjected to forces from the vertex model work function and the polarity-directed active forces. **b,** Dynamic phase diagram of the active vertex model, color-coded with cell-cell re-arrangement rate, with a dashed-line separating the solid and the flowing regimes. **c,** Simulations of the active vertex model in the solid (**i**) and flowing (**iii**) regimes, and in transition regime between the two (**ii**). The top row shows a single snapshot with polarity denoted by red arrows. The second row shows the angular position of the rotation axis through time in the (*θ, ϕ*) polar plot. The third row shows the magnitude of the angular velocity and the angular change of the rotational axis. **d-e,** The solid body rotation fraction of the surface velocity *f*_rot_ (see Methods) in simulations of the solid and flowing regimes of active vertex model. **f,** The rotation fraction *f*_rot_ in experiments indicates that pancreas spheres rotate as an active solid.

The cell polarity 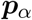 directs the traction force exerted by the cell *α* on the surrounding matrix. The time evolution of cell polarity follows the dynamics:

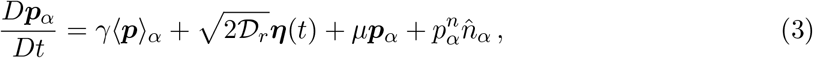

where *D/Dt* denotes a co-rotational time derivative (see SI Sec. 3). The first term on the r.h.s. of the equation (3) accounts for the alignment of cell polarity with the average polarity of its *M_α_* nearest neighbors 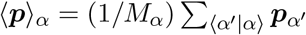 with a rate *γ*. The second term accounts for a rotational noise with a diffusion coefficient 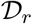. The polarity noise 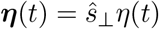 is perpendicular to both cell polarity and the normal vector at the cell center 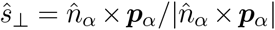. The polarity noise magnitude *η* is a Gaussian variable with mean 0 and variance 1. The third term is used to impose 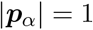 at each time through a Lagrange multiplier 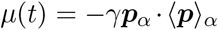. With the last term we ensure that the polarity remains in the tangent plane of the sphere by adding a normal component with the magnitude 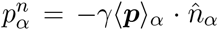. In this study, we initialize the tissue with the force balanced configuration of a Voronoi diagram construction of *N* randomly distributed cell centers on a sphere of radius *R_s_* = (*NA*_0_*/*4*π*)^1*/*2^, and we initialize the direction angle of cell polarity from a uniform random distribution (see SI Sec. 3).

We first explore the vertex model dynamics on a sphere in the absence of noise by setting 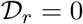. We identify two distinct dynamic regimes; the solid and the flowing regimes. In the solid regime cells do not rearrange. At finite alignment rate *γ* > 0 the polarity vectors are globally aligned (Fig. 2c(i), SI Video 6) and the sphere rotates as a solid body around a constant axis, consistent with the experimentally-observed rotational motion (Fig. 1f(i)). In the flowing regime cells continuously rearrange and the sphere rotation speed is significantly lower than in the solid regime (Fig. 2c(iii)). We identify these regimes in a dynamic phase diagram (Fig. 2b) by measuring the cell rearrangement rate *R*. The flowing regime appears only when the traction force magnitude *F* is greater than the threshold value *F_c_* which increases with the alignment rate *γ*. Near the transition, the sphere dynamics is erratic with sudden bursts of cell rearrangements occasionally punctuating the otherwise solid-like rotating state, leading to changes in the orientation of the rotation axis (Fig. 1c(ii), SI Video 7). This behaviour is reminiscent of the yielding transition [26], and the sphere dynamics is consistent with the rotation axis drift found in experiments (Fig. 1g(ii)).

A key question is which dynamic regimes do rotating pancreas spheres exhibit. Due to the high uncertainty to identify cell rearrangements in the experimental data, a reliable measurement of the rearrangement rate *R* was not feasible. Instead, we address this question by decomposing the surface velocity field **V**(*θ, ϕ*) in spherical harmonic modes. We then define the solid body rotation fraction quantity *f*_rot_ that captures the solid body rotation component of the full surface velocity field (Fig. 2d, see SI Sec. 2.4). In our simulations in the solid regime, the surface velocity **V**(*θ, ϕ*) has a high fraction *f*_rot_ > 0.8 of *l* = 1 (rotational) mode, compared to the flowing regime with fraction *f*_rot_ ≈ 0.3 of this mode. In experiments, the quantified surface velocity **V**(*θ, ϕ*) of pancreas spheres has a fraction of the rotation mode of *f*_rot_ ≳ 0.8, consistent with signatures of solid rotation in the active vertex model (Fig. 2e, SI Fig. 3). Altogether, the quantitative comparisons between experiments and simulations indicates that pancreas spheres rotate as active solids, with some spheres positioned near the yielding transition regime that shows drift in the rotation axis.

### Polarity alignment and traction force control cell shape patterns

To further investigate how polarity alignment and traction force govern different aspects of sphere rotational dynamics, we performed single cell segmentation (Fig. 3a, see SI Sec. 2.5) and quantified two key cell shape parameters: (i) norm of the cell elongation tensor 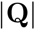 and (ii) cell elongation orientation angle *β* with respect to the azimuth in the direction of motion (Fig. 3(b)). Surprisingly, in experiments, we found that the cell shape orientation is nearly aligned system-wide (Fig. 3c) with the orientation angle distribution *P* (*β*) showing a pronounced peak at either *β* ≈ 45° or *β* ≈ 135° (Fig. 3d). In the active vertex model, when polarity alignment is turned off, the random traction force will still result in a net torque that generates a rotation. However, in this case, the rotation speed is slower and the cell shape orientation does not show a preferred angle *β* (Fig. 3e) and *P* (*β*) shows a flat distribution. This distinction from experiments indicates that polarity alignment is required to generate the observed rotational dynamics in pancreas spheres.

**Figure 3:**
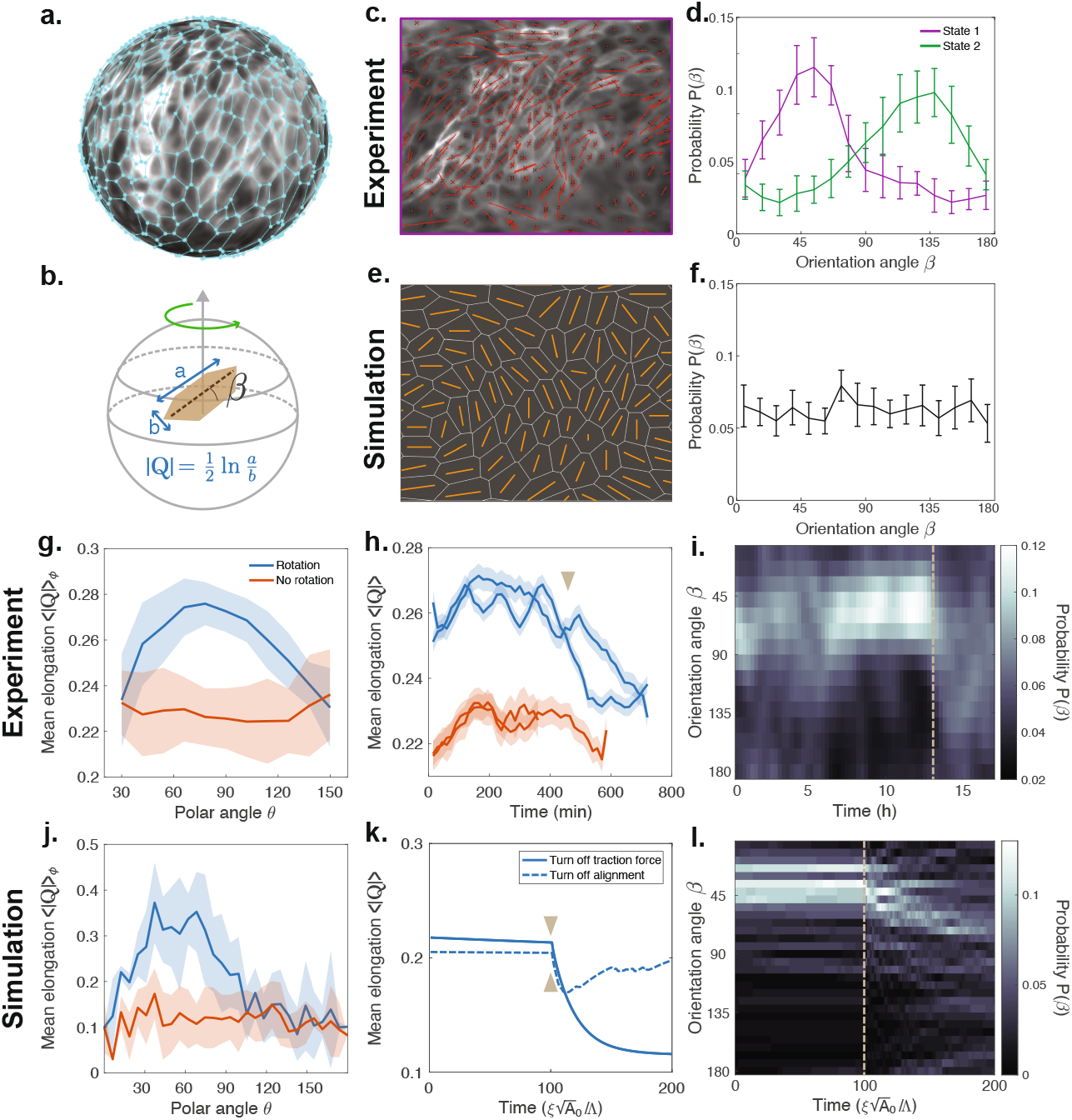
Polarity alignment and traction force control cell shape patterns. **a,** Cell segmentation performed on the projection of pancreas sphere. **b,** Schematic defining cell shape elongation parameter 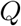 and cell shape orientation angle *β* (defined with respect to azimuth). **c,e,** A Mercator projection showing the cell elongation pattern in experiments (**c**) and active vertex model (without polarity) (**e**). **d,** The cell shape orientation distribution *P* (*β*) in two different experiment states, showing a unimodal distribution with a peak at either 45° or 135°. **f,** The cell shape orientation distribution *P* (*β*) in active vertex model (without polarity) shows a uniform distribution. **g-i,** Cell shape dynamics in experiments during rotation arrest. Mean cell elongation 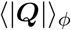 (azimuthal average) as a function of polar angle *θ* before (blue curve) and after (orange curve) rotation arrest (**g**). Mean cell elongation 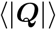 (whole system average) as a function of time for 2 spheres that undergo rotation arrest (blue curves, brown arrow denotes start of arrest) and 2 spheres that do not rotate (orange curves) (**h**). Probability-time kymograph of cell orientation angle *β* for a sphere that undergoes rotation arrest (**i**, brown dotted line denotes start of arrest). **g-i,** The corresponding cell shape dynamics in the active vertex model during rotation arrest. In (**k**), we compare the mean cell elongation 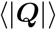 (whole system average) when either traction force (solid blue curve) or polarity (dotted blue curve) is turned off. In (**j**) and (**l**), we show the plots for the case when traction force is turned off.

To investigate the role of traction forces in sphere dynamics, we focus on cell shape dynamics during rotation arrest. We found that during rotation, the cell shape elongation (as quantified by the mean cell elongation 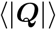, see SI Sec. 2.5) shows a maximum near the equator (polar angle *θ* = 90° ), with cells at both poles being more isotropic (Fig. 3g, blue line). In contrast, cell elongation in non-rotating spheres is roughly spatially homogeneous (Fig. 3g, orange line). Interestingly, we found that as spheres stop rotating, the shape of cells becomes more isotropic, reaching a level that is comparable to cells in non-rotating spheres (Fig. 3h). Within the active vertex model, a sphere can stop rotating in two ways: (i) by turning off the traction force or (ii) by turning off the alignment rate. We reason that the experimental observation corresponds to case (i), since turning off alignment while keeping traction force constant would put the system in a mechanically frustrated state, where polarity vectors are misaligned and cells would become more stretched. Indeed, by performing the respective simulations (Fig. 3k), we found that turning off traction force recapitulates the reduction in cell shape elongation as observed during rotation arrest in experiments. Consistent with experiments, the mean elongation of cells near the equator is higher in rotating spheres, compared to non-rotating spheres (Fig. 3j). Furthermore, we found that the cell orientation *β* exhibits a unimodal distribution during rotation (with a peak ≈ 45°), which subsequently homogenizes after rotation arrest (Fig. 3e). This behavior is similarly observed in our simulation with traction force turned off (Fig. 3h).

### Chiral symmetry breaking in the cell shape orientation field

Cell morphology patterning is known to encode information about tissue mechanics during development [28] and apoptosis [6]. In the absence of external signaling, as in the case of a rotating sphere, we do not expect the cells to be oriented in any specific pattern. Remarkably though, we found that the cell shape orientation *β* in rotating pancreas spheres shows a preference to be either ≈ 45° or ≈ 135° for a sustained period of time (Fig. 4a). This is clearly reflected in the unimodal orientation distribution *P* (*β*), which in non-rotating sphere is distinctively uniform (SI Fig. 5). The fact that these cell arrangements are mirror images (and thus cannot be superimposed on one another, Fig. 4b(i)) imply that chiral symmetry has been spontaneously broken. Careful reasoning reveals that up-down symmetry is also broken (SI Fig. 4). When we computed the mean orientation angle 〈*β*〉_*ϕ*_ (averaged azimuthally), we found that the cell orientation is roughly independent of the polar angle *θ* (Fig. 4c, SI Fig. 6).

**Figure 4:**
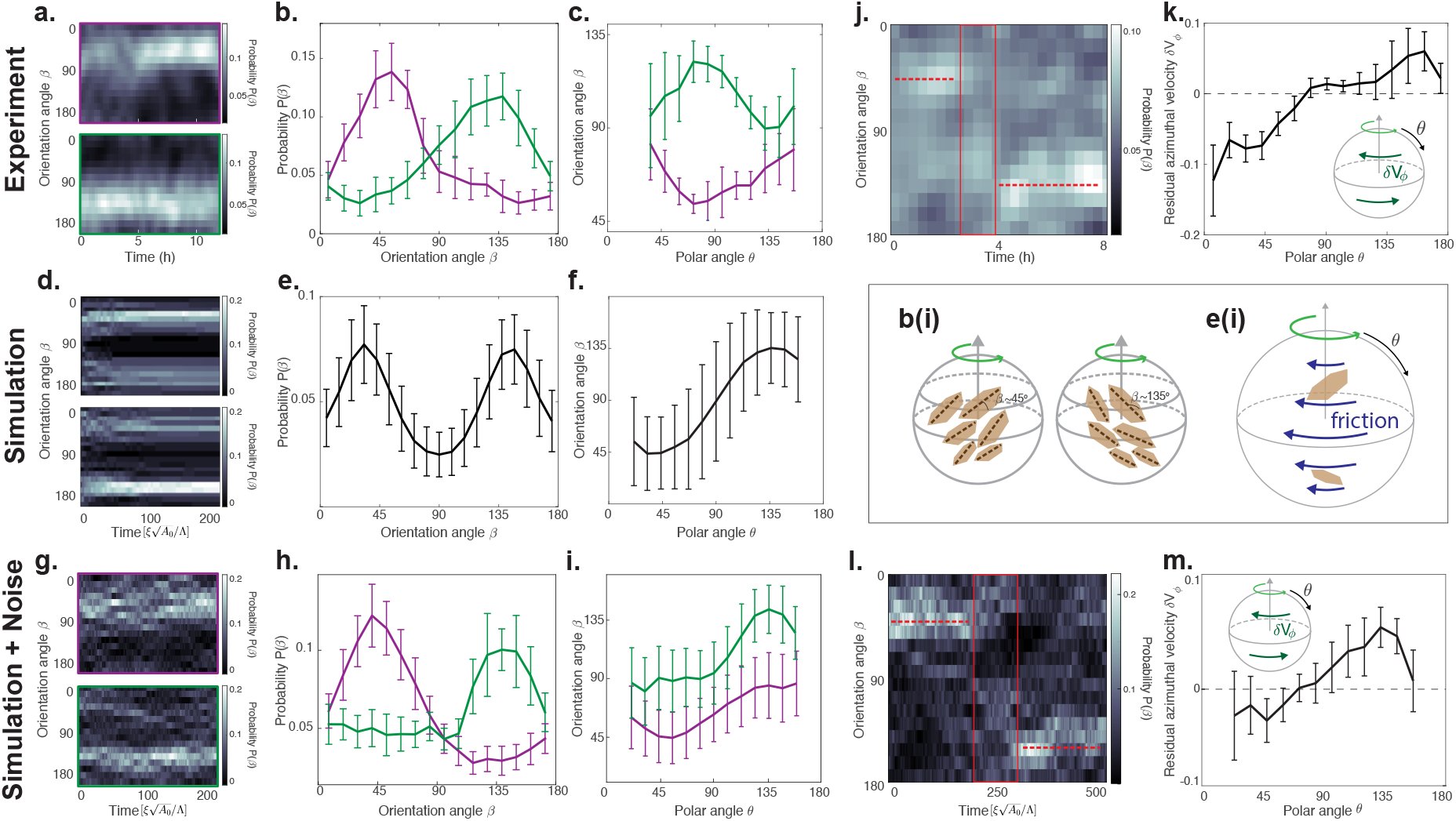
Chiral symmetry breaking in the cell shape orientation field. **a,** Probability-time kymograph of the cell orientation *β* for two different pancreas spheres undergoing persistent rotation. **b,** The cell orientation probability distribution *P* (*β*) for the two spheres in (**a**) shows unimodal distributions with peaks at around 45° (purple) and 135° (green). **c,** Azimuthally-averaged cell orientation angle 〈*β*〉_*ϕ*_ as a function of polar angle *θ* for the two spheres in (**a**). Note that *β* remains less (more) than 90° for the purple (green) state for the entire sphere, as illustrated in the schematic (**b**(i)). **d-f,** Analogous cell orientation analysis performed on simulations (without noise). *P* (*β*) shows bimodal distribution with peaks at both 45° and 135° (**e**). The polar dependence of 〈*β*〉_*ϕ*_ (**f**) indicates that cell orientation is symmetric about the equator, as illustrated in the schematic (**e**(i)). **g-i,** Analogous cell orientation analysis performed on simulations (with noise). **j,l,** Probability-time kymographs of cell orientation *β* showing a switch of preferred angle from 45° to 135° in experiment (**j**) and theory (**l**). **k,m,** The residual azimuthal velocity *δ***V**_*ϕ*_ (see Method) as a function of polar angle *θ* for experiment (**k**) and simulation (**m**). Insets show the directionality of *δ***V**_*ϕ*_ with respect to angular velocity.

**Figure 5:**
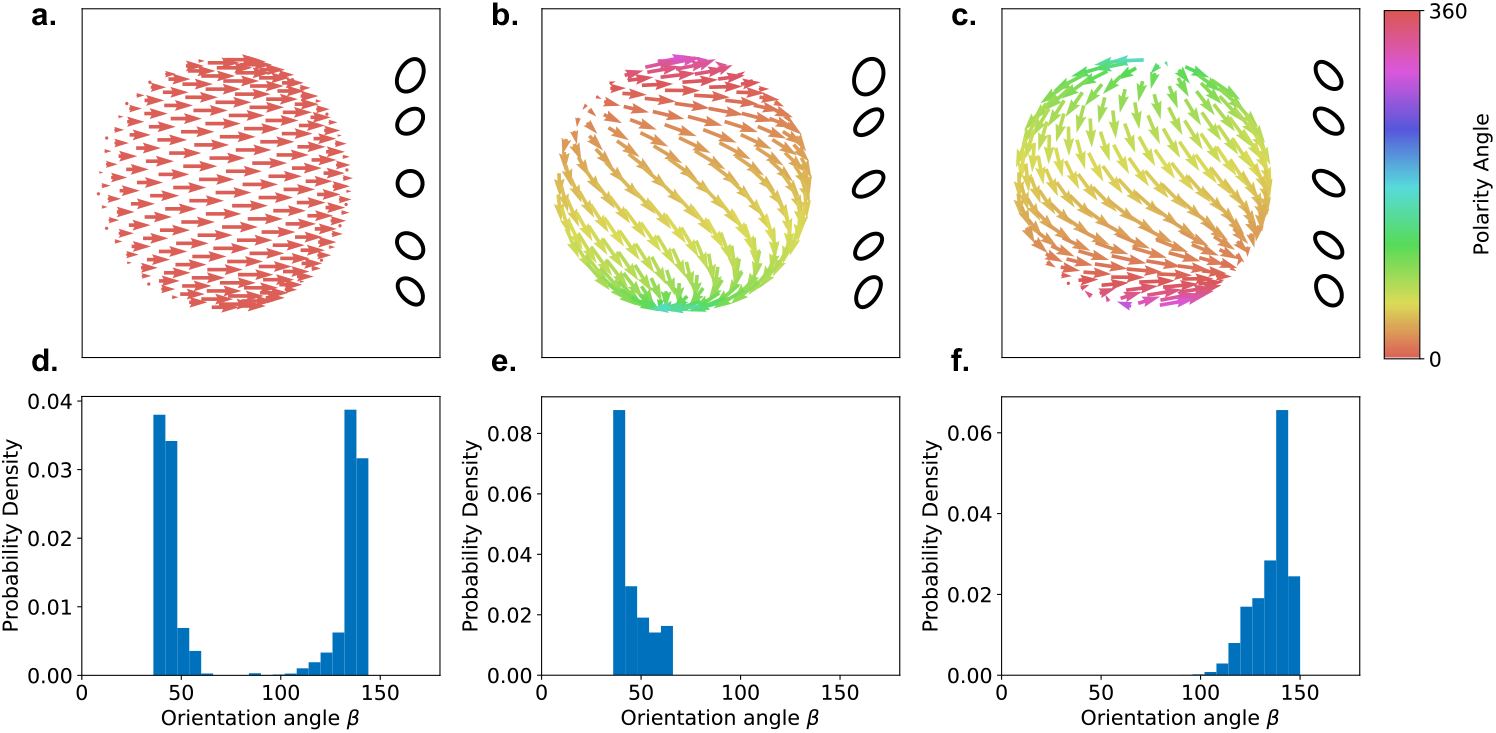
Topological defects in the polarity field underlie chiral symmetry breaking. **a-c,** Axisymmetric solutions to 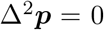 with 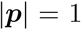 of the polarity field 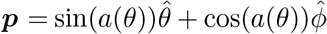 parameterized by an angle *a(θ)*, see SI Sec 5. Up-down symmetric solution with two vortices at the poles and angular profile *a*(*θ*) = 0 (**a**), a vortex near the north pole and an aster near the south pole, 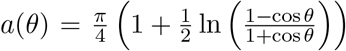 (**b**), and the mirror image with respect to the equator, 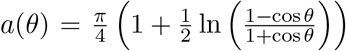 (**c**). The ellipses indicate cell shape and cell elongation at different polar angles *θ*. **d-f,** Inferred cell elongation orientation distribution *P* (*β*) corresponding to the three polarity patterns in (**a-c**).

To investigate the nature of this broken chiral symmetry, we turned to the active vertex model. Surprisingly, we found that in simulations, the orientation distribution *P* (*β*) is bimodal, with peaks at both ≈ 45° *and* ≈ 135° (Fig. 4d-e). One explanation for this co-existence of both orientation angles comes from the spherical geometry of the system. In solid sphere rotation, cells near the equator move faster than those near the pole, thus generating a gradient of shear stress induced by friction along the polar direction *θ* (Fig. 4e(i)). This could in principle result in shear alignment of the cell elongation axis, which has been shown to behave as a nematic director. Such a mechanism would predict a symmetric cell orientation profile along the equator, with cell orientation predominantly taking the value of ≈ 45° at the ‘northern’ hemisphere (*θ* < 90°) and ≈ 135° at the ‘southern’ hemisphere (*θ* > 90°) (Fig. 4e(i)). Indeed, the measured orientation angle *β* from simulations shows the expected *θ* dependence (Fig. 4f).

While the spherical geometry provides an explanation for the existence of two preferred angles, it doesn’t explain how the symmetry is broken in experiments. The symmetric cell shape orientation profile in simulations (Fig. 4d-f) involves an abrupt change in cell elongation orientation which generically costs elastic energy. We posit that stochastic noise in experiments could help bias cell orientation to minimize this large bending energy. By introducing a noise term into the polarity equation of active vertex model (see SI Sec. 3), we found that the symmetry does indeed get broken and the orientation angle distribution *P* (*β*) becomes unimodal with peaks at either ≈ 45° or ≈ 135° (Fig. 4h-i). This result suggests that the cell orientation field is mechanically poised in a bistable state, and can thus be pushed into either state through stochastic noise. Indeed, in longer simulations, we found that the cell orientation *β* can switch between the two states (Fig. 4l). During this window of switching, we detected a counter-clockwise twist in the residual azimuthal velocity *δ***V**_*ϕ*_, which generates the required shear gradient to flip the preferred orientation *β* from 45° to 135° (Fig. 4m). Such a switch is also observed in experiments (Fig. 4j), with the expected residual azimuthal velocity profile (Fig. 4k). Furthermore, in both experiment and simulation, we observed the opposite switch in preferred angle from 135° to 45° (SI Fig. 7). Altogether, these results show how geometrical confinement and stochastic noise in multicellular systems can lead to the emergence of chirality in tissue patterning.

To investigate the underlying symmetry breaking mechanism, we turned to a simplified continuum model of the sphere as an elastic shell with a polarity field generating traction forces (SI Sec. 5). We determine polarity fields that solve the Laplace equation 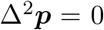 with 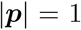, the continuum equivalent of neighbour alignment. We consider solutions that are symmetric about the axis of rotation (SI Fig. 9). The sphere is described as an elastic spherical shell that is not deforming radially and obeys the force balance equation 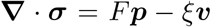, where ***σ*** is the elastic stress tensor, ***p*** and ***v*** are the polarity and sphere surface velocity respectively, *F* is traction force and *ξ* the friction coefficient. The velocity is found from torque balance between the traction forces and frictional forces, assuming solid body rotation (SI Sec. 5.2). Solving for the polarity field, we find a family of solutions that can be specified by two parameters (see SI Sec. 5.1, SI Fig. 9). By varying the two parameters, we can select up-down symmetric solutions (Fig. 5a) and solutions with broken up-down symmetry (Fig. 5b-c).

In order to relate these solutions to patterns of orientation of cell elongation, we calculate from the given polarity field an angle that corresponds to the tilt angle *β* of cells. It is defined via the orientation of shear deformations on the surface (see SI Sec. 5.3). Importantly, considering the up-down symmetric polarity field containing two vortices at the poles, the cell orientation distribution *P* (*β*) exhibits a bimodal shape (with peaks at ≈ 45° *and* ≈ 135°, Fig. 5d).Note that this pattern is not chiral. In contrast, considering polarity field with broken up-down symmetry, the distribution *P* (*β*) exhibits a unimodal shape, with a single peak at either ≈ 45° (Fig. 5e) or ≈ 135° (Fig. 5f). These patterns with broken up-down symmetry are also chiral, with opposite handedness. Altogether, these results suggest that the positioning and asymmetry of the topological defects of the polarity field govern the chiral symmetry breaking observed in the rotating cell orientation field. Further analysis of the chiral states in the noisy active vertex model reveals the signature of the broken up-down symmetry in the polarity field (SI Fig. 8), consistent with the continuum model.

## Discussion

In this work, we show how coupling of cells in 3 dimensions can modulate the collective cell behavior and the emergent mechanics of multicellular systems. Specifically, we demonstrate how an active vertex model can account for distinct rotational dynamics including persistent rotation, drift in angular velocity and rotation arrest in pancreas spheres. The mechanisms we uncover are likely to be conserved in rotating spheres derived from other cell types [21, 22, 29], with potential implications for cell migration on complex 3D curved geometries (e.g. polarized tissue flow in the *Drosophila* embryo [30]).

Surprisingly, we find that the topological constraints on a spherical epithelium can poise the system in a bistable state in which noise can spontaneously break chiral symmetry. Results from our continuum model demonstrate that an asymmetry of topological defects in the polarity field underlies the two coexisting chiral states. This process of generating tissue chirality through topological constraints is distinct from previously reported mechanisms which arise from either rotation of nodal cilia [31], chiral cell polarity [32], or a chiral phase that arises generically from a planar polarized epithelial system [33]. More generally, our results suggest an alternative route where a localized biological signal could drive symmetry breaking processes in tissues by controlling the types and locations of topological defects in cell polarity.

## Acknowledgments

T.H.T. and M.F.S. acknowledge support from the Center for Systems Biology Dresden as ELBE Postdoctoral Fellow. A.A. acknowledges support from the Federal Ministry of Education and Research (Bundesministerium fuür Bildung und Forschung, BMBF) under project 031L0160. C.D. acknowledges the support of a postdoctoral fellowship from the LabEx “Who Am I?” (ANR-11-LABX-0071) and the Université Paris Cité IdEx (ANR-18-IDEX-0001) funded by the French Government through its “Investments for the Future” program. We are grateful to the Light Microscopy Facility and the Biomedical Services at the MPI-CBG for the support in all the imaging and maintenance of the mice, respectively.

## Supplementary Information

### 1 Experimental setup

#### 1.1 Animal lines and permits

All experiments were performed in accordance with the German Animal Welfare Legislation (“Tierschutzgesetz”) after approval by the federal state authority Landesdirektion Sachsen (license DD24.1-5131/451/8). Mice were kept in standardized specific-pathogen-free (SPF) conditions at the Biomedical Services Facility (BMS) of Max Planck Institute of Molecular Cell Biology and Genetics. Genetically modified mouse lines ‘Myh14<tm3.1Rsad>/Mmjax’ (Jackson Laboratory stock number 023449) and ‘Gt(ROSA)-26Sortm4-(ACTB-tdTomato-EGFP)-Luo(C57Bl/6NCrl;Crl)’ [1] were used to follow the Myosin and cell membrane patterns, respectively.

#### 1.2 Preparation of organoid culture

To generate mouse pancreas spheres, we follow the protocol developed in [2] with some modifications. Briefly, we dissect the developing dorsal pancreas and harvest progenitors cells from mice at embryonic day (E) 13.5. We obtain single cells by a combination of chemical and mechanical dissociation: we incubate with TrypLE (GIBCO-BRL, Invitrogen 12604-013) at 37° for 7 min in conical wells of 60-well mini-trays (nunc, 439225), transfer the buds into a well with fresh DMEM (Gibco 10565-018) and perform the mechanical dissociation by aspirating and ejecting the bud into 10 μl pipette tips until complete dissociation under microscopic control. Then, we pool the cells from 4 embryos into an Eppendorf tube in order to minimize differences due to individual processing. We dilute the cell suspension to a final concentration of 75% Matrigel (Corning Costar 356231) on ice to prevent Matrigel polimerization during the mixing process. We prepare 5 μl drops in a pre-treated chamber for imaging (see 3D imaging below) and incubate the chamber at 37°C for 30 min for the polymerization of the Matrigel. The wells are then filled with 250 μl of sphere medium (DMEM/F12 + Glutamax (Gibco 10565-018), 1% Pen/Strep (Gibco 15140-122), 10 μM Y-27632 (SIGMA), 10% Y0503-IMG B27 x 50 (ThermoFisher 17504-044), 64 ng/ml FGF2 (R&D Systems 450-33-10UG) [2]) and incubated in a humidified environment containing 5% CO_2_ and 95% air at 37°C. We replace the medium to a new sphere medium without the ROCK inhibitor (Y-27632) after 2 days and again after 4 days and image at day 5.

#### 1.3 3D live imaging of pancreas spheres

Sphere samples are placed in Fluorinated ethylene propylene (FEP) bottom V-shaped chambers from Viventis previously treated as follows: we fill each well with distilled water, then replace it with 70% ethanol after 5 minutes; we remove the ethanol and rinse once with distilled water and let it dry under the hood for 20 minutes and finally under UV for 20 minutes. After 5 days of sphere culture, time lapse 3D imaging is initiated using the LS1 Live light sheet microscope system from Viventis. The parameters used are the following: dual illumination, 2.2 μm beam; Laser 488/561, exposure 50 ms and detection objective Nikon 25X NA 1.1.

### 2 Data analysis

#### 2.1 Sphere surface segmentation and projection

To obtain the projection of the surface fluorescence intensity of the pancreas spheres, we first obtain the segmentation of the sphere apical (inner) surface. This is done using the following steps: First, we train a pixel classification model using ilastik [3] to distinguish between ‘tissue pixels’ and ‘background pixels’ in our 3D microscopy *z*-stack images. Using this model, we create a 3D volume mask that identifies the voxels that represent the tissue volume. Next, we use the MATLAB function bwboundaries to identify the inner surface of the sphere. By performing this step for the entire *z*-stack, we obtain the point cloud 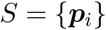 that defines the inner surface of the 3D sphere. Using this point cloud, we extract the apical surface fluoresence intensity *I*(*x, y, z*). To simplify downstream analysis, we make the assumption that all pancreas spheres are roughly spherical, and project the intensity onto a sphere to obtain *I*(*θ, ϕ*), using the transformation *θ* = arctan 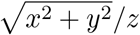 and *ϕ* = arctan *y/x*.

#### 2.2 Reconstruction of the 3D tissue flow field

To reconstruct the full 3D tissue flow field, we first perform 2D particle image velocimetry (PIV) on three different projections: the equatorial projection, the north pole projection and the south pole projection. The equatorial projection on a 2D plane is defined by *x*′ = *ϕ* and *y*′ = *ϕ* (here, we use prime notation to represent the projection coordinates, while the non-prime notation represents real space coordinates). The north and south pole projections are defined as *ϕ*′ = ±*ϕ* and *r*′ = (*θ, π* − *θ*), respectively, where *θ* = [0, *π*/2] for north pole projection and *θ* = [*π/*2*, π*] for south pole projection. Next, we perform 2D PIV on the three projections separately, using the application PIVlab in MATLAB, developed by [4]. To reconstruct the full 3D flow fields, we stitch the PIV fields from the three different projections, using the north projection for *θ* = [0, *θ_c_*], the equatorial projection for *θ* = [*θ_c_, π* − *θ_c_*], and the south projection *θ* = [*π* − *θ_c_, π*]. The rationale for this stitching scheme is to use regions of PIV fields from different projections that minimize distortion. *θ_c_* ≈ 1 radian is determined by considering *θ* dependence of the distortion for each projection. To ensure smoothness of the stitched velocity field, we further define an overlap region of ±Δ*θ* where the velocity fields from two different projections are averaged.

#### 2.3 Computation of the angular velocity Ω(t)

This section explains how we obtain the angular velocity **Ω**(*t*) and the residual velocity from the motion of vertex model spheroids and the experimental spheres. We denote the position of each vertex of the vertex model at time *t* by ***r**_i_*(*t*), and the vertex velocity is estimated as

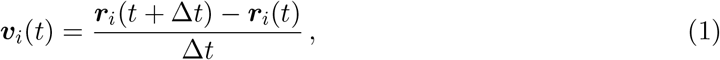

where Δ*t* is the time interval between two subsequent frames. On the other hand, for the experimental organoids, we find the positions and velocities on the surface using PIV (see SI Sec. 2.2). In the following we use the term ‘*points*’ to refer to both vertices of the vertex model epithelium and also the tracked points from PIV analysis of the spheres.

We first define the average position ***R***(*t*) and average velocity ***V*** (*t*) of the tracked points as

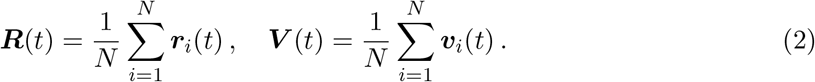

In full generality, the velocity of point *i* can be decomposed as

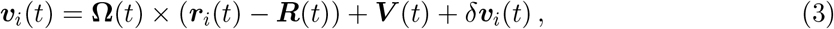

where **Ω**(*t*) is an angular velocity that will be computed in the following and which is defined such that *δ**v**_i_*(*t*) = **0** if the *N* points are placed on a solid body with center of mass ***R***(*t*), moving at velocity ***V*** (*t*) and rotating with angular velocity **Ω**(*t*).

In order to compute **Ω**(*t*), we define the angular momentum **Γ**_*i*_(*t*) of point *i* as

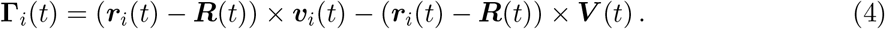

Inserting this definition into Eq. (3) yields

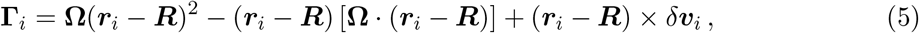

where the time dependence has been omitted for simplicity. Equation (5) can then be rewritten in matrix form as

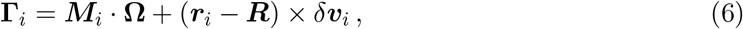

where we have introduced the moment of inertia tensor ***M**_i_* of point *i*

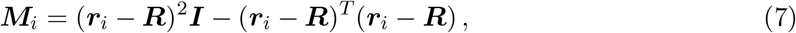

with ***I*** the identity tensor. The average angular momentum is obtained as

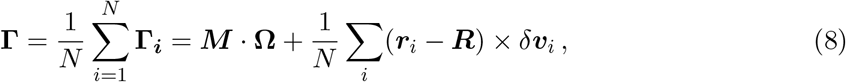

where 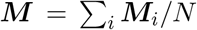 is the average moment of inertia tensor. In the case of a solid-body rotation, *δ**v**_i_* such that the angular velocity is simply obtained from Eq. (8) as

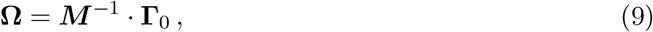

with 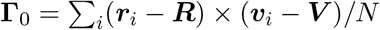.

The residual motion *δ**v**_i_*(*t*) is then obtained as

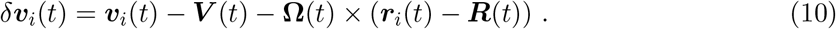

Note that the computation of the solid-body angular velocity **Ω** requires to invert the moment of inertia tensor ***M***, and thus requires *N* > 2.

#### 2.4 Vector Spherical Harmonics Decomposition and definition of solid body rotation fraction *f_rot_*

The velocity field **V**(*θ, ϕ*) on a sphere can be decomposed into real vector spherical harmonics (SH) modes as

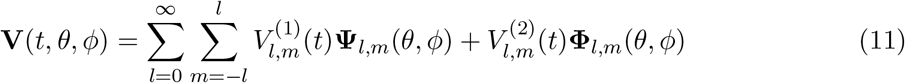

where, for each mode (*l, m*), 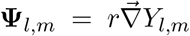 and 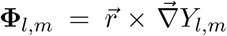 are the curl-less and divergence-less components of the vector spherical harmonics, respectively, and *Y_l,m_* are the real scalar spherical harmonics defined as:

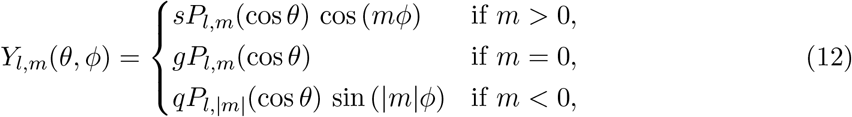

where 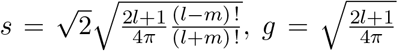 and 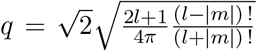 are normalization factors. The components of the velocity field are then defined as

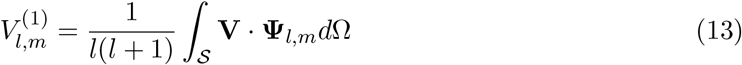

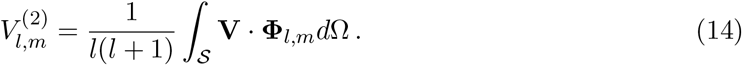

The fraction of solid body rotation in the surface velocity **V**(*θ, ϕ*) is defined as

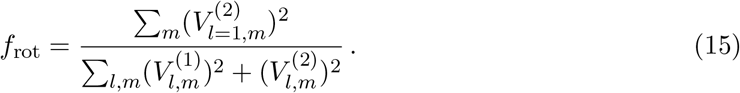

#### 2.5 Single cell segmentation and quantification

To obtain the single cell shape quantification, we first perform segmentation of the pole projections using the automated segmentation package EPySeg [5]. Once the cell boundaries are identified, the MATLAB function regionprops is used to find regions corresponding to single cells and calculate their geometrical properties. Each cell region is fitted with an ellipse that has the same normalized second moments as the region. The cell elongation is defined as 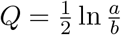 where *a* and *b* are the lengths of the major and minor axes, respectively. The cell orientation *β* is defined as the angle between the major axis and the local latitude (i.e., the 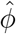 direction).

## 3 Active vertex model on sphere

We consider a two-dimensional vertex model with cell polarity on a rigid sphere. Cells on the sphere are represented as polygons that are outlined by straight edges connecting vertices [6]. We consider a polygonal cell network consisting of *N*_c_ cells labelled *α*, *N*_v_ vertices labelled *m* and *N*_b_ straight bonds between connected vertices labelled 〈*mn*〉 where *m* and *n* are the vertices they connect. Each cell is characterized in terms of its area *A^α^*, perimeter *L^α^* and polarity 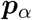.

In our vertex model, the forces stemming from the mechanical properties of individual cells and from the interactions between them originate from the vertex model work function

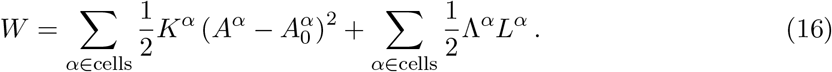

The first term describes an area elasticity contribution, with 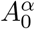 the preferred cell area and *K^α^* the area stiffness. The second term describes a contribution due to the tension of network bonds, where Λ^*α*^ is the perimeter tension magnitude, and *L^α^* denotes cell perimeter [6, 7]. Here, for simplicity, we consider the case where the cell parameters are identical for all cells in the tissue: 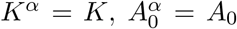 and Λ^*α*^ = Λ. Due to the spherical geometry, polygonal cells are not co-planar and a unique definition of cell area requires a suitable triangulation. Here, we define the cell center ***X***_c_ as the average position of vertices, and triangulate each cell by connecting the cell center to consecutive cell vertices ***X**_i_* and ***X**_i_*_+1_ in counter-clockwise order (see Figure below). Hence, for a given cell, we obtain the cell area and perimeter as

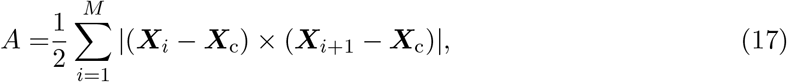

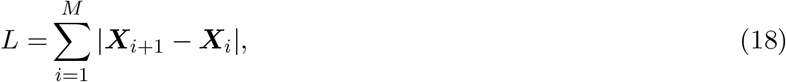

where *M* is the number of vertices of the cell, and with ***X**_M_*_+1_ ≡ ***X***_1_.

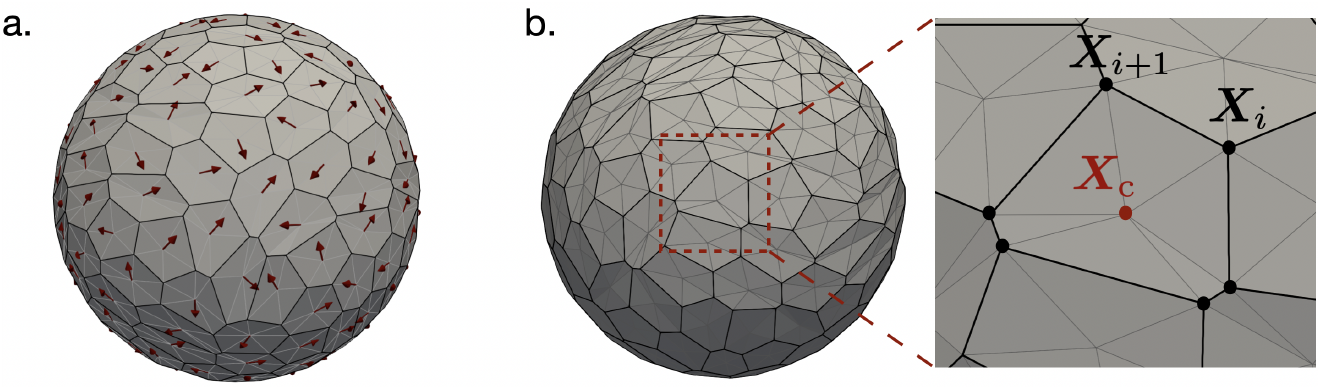

**a.** Example of an initial condition of the tissue configuration and of the cell polarities. **b.** Tissue triangulation and definition of geometric quantities.

At each vertex *m* located at position ***X**_m_*, force balance reads

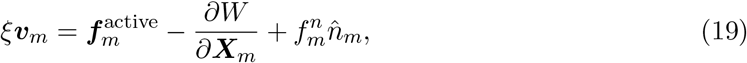

where 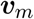 is the velocity of vertex *m*, and *ξ* is the friction coefficient with the external environment. The traction forces 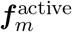 are generated from active processes and are defined on each vertex as

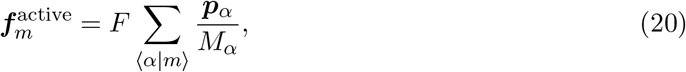

where *F* denotes the traction force magnitude of a cell and the sum is over the (three) cells that share vertex *m*, and *M_α_* is the number of vertices of cell *α*. We consider a non-deforming spherical geometry by setting 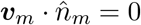, which determines the normal force as

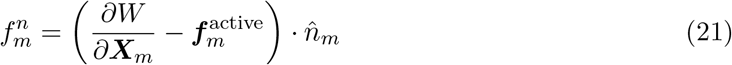

where 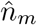 is the unit normal vector to the sphere at vertex *m*.

We consider a polarity field 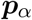 in the tangent plane that directs the traction force exerted by the cell *α* on the surrounding matrix. The time evolution of the polarity direction follows the dynamics

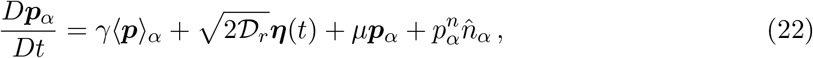

where *D/Dt* denotes a co-rotational time derivative (see SI Sec. 4). The first term on the right-hand side of the Eq. (22) accounts for the alignment of the polarity of cell *α* with the polarity of its nearest neighbors with a rate *γ* with the definition

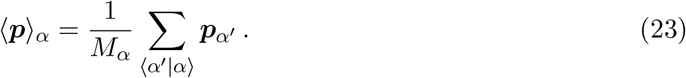

The second term in Eq. (22) accounts for the rotational noise with a diffusion coefficient 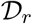. The white Gaussian noise 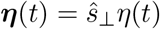 is perpendicular to both cell polarity and the outward normal vector to the cell 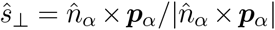 and has a random magnitude *η* that is defined as a Gaussian variable with mean 0 and variance 1. The Lagrange multiplier 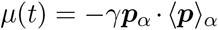 imposes 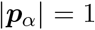 at each time. With the last term we ensure that the polarity remains in the tangent plane by adding a normal component with the magnitude

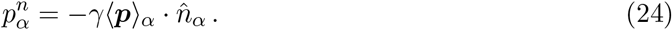

The dimensionless model parameters are defined as 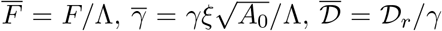, and 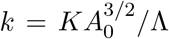 We fix *k* = 10, which ensures mechanical stability of the cellular network as we vary the remaining parameters. In this study, we initialize the tissue by a force balanced configuration of a Voronoi diagram construction of *N* randomly distributed cell centers on a sphere of radius *R_s_* = (*NA*_0_*/*4*π*)^1*/*2^, we initialize the cell’s polarity angle form a uniform random distribution.

## 4 Implementation of the co-rotational time derivative

The cell polarities are transported with the cells through the co-rotational time derivative in Eq. (22). This equation can be written for a time step Δ*t* in a discrete form

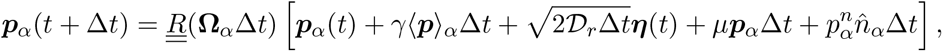

where the three dimensional rotation matrix 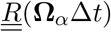 rotates the polarity vector with the cell and is constructed by extracting the solid body angular velocity **Ω**_*α*_ of cell *α* from the velocity of its vertices

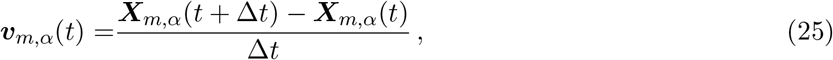

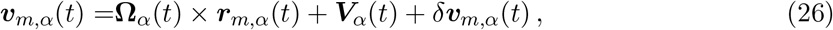

where ***X**_m,α_*(*t*) is the position of each vertex *m* that belongs to cell *α*, and ***r**_m,α_*(*t*) = ***X**_m,α_*(*t*) − ***X**_α,c_*, and ***X**_α, c_* denotes the cell center position. The decomposition of velocity and the determination of the cell angular velocity follows the method described in SI Sec. 2.3.

## 5 Continuum model of cell polarity

### 5.1 Polarity dynamics

We model the cell polarity by a polar field 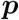 on the surface of the sphere of radius *r*. In the continuum limit of neighbour alignment, the polarity evolves according to

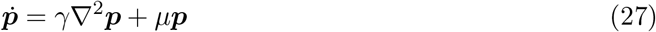

where *γ* is the alignment rate and *μ* is a Lagrange multiplier to impose 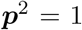. We solve for axisymmetric steady-state solutions of polarity, expressed as

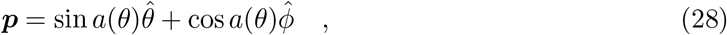

imposing that polarity is a unit vector described by the angle *a*(*θ*) with respect to the 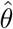 direction.

The Laplacian of polarity is given by

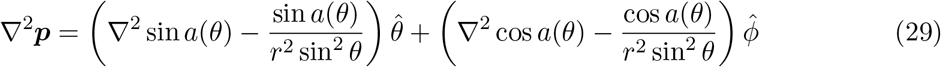

and thus the polarity evolves according to

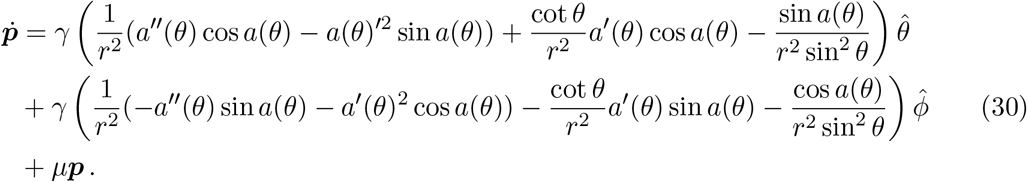

Imposing the constraint 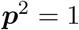 we finally obtain

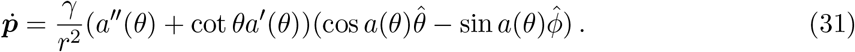

The corresponding dynamical equation for the polarity angle *a(θ)* reads

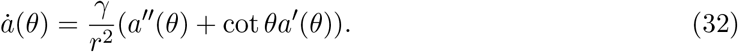

It has steady state solutions (SI Fig. 9) of the form

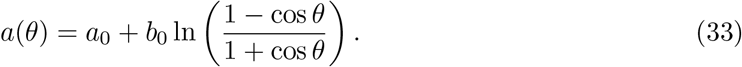

This solution contains two defects in the polarity field located at opposite poles of the sphere. Singularities appearing in the polarity field at *θ* = 0 and *θ* = *π* are regularised by the presence of defect cores of size *s*, that are equal or larger than the linear cell size.

Up-down mirror symmetric defect configurations correspond to *a*_0_ = 0 (see SI Fig. 9) which produces a symmetric divergence of the polarity field with respect to *θ* → *π* − *θ* on the sphere

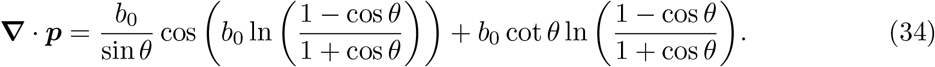

Up-down anti-symmetric defect configurations correspond to *a*_0_ ≠ *b*_0_ = 0, which produce an anti-symmetric divergence of the polarity field on the sphere

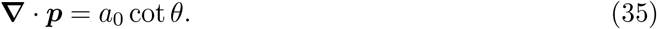

### 5.2 Elastic shell and displacement field

We describe the tissue as a spherical elastic shell with shear modulus *G* and bulk modulus *K*, and stress tensor ***σ***. Using the steady state polarity field, which generates a traction force 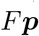 on the surface of the sphere, we calculate the displacement field **u**, on the sphere’s surface to find the predicted cell tilt angle *β* of local elongation. The sphere rotates like a solid body with angular velocity *ω* due to torque balance with frictional forces from the surrounding ECM, giving a surface velocity of **v** = *ω* × **x**. Force balance is then given by

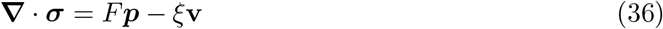

where *ξ* is the friction coefficient.

We obtain the angular velocity through torque balance

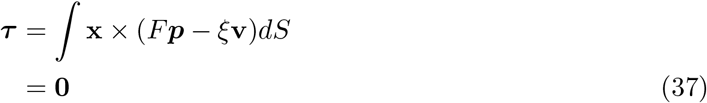

To effectively solve these equations, we decompose the polarity, displacement and velocity fields into vector spherical harmonics as

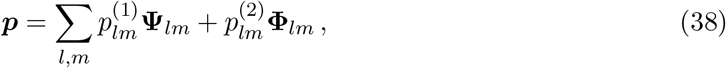

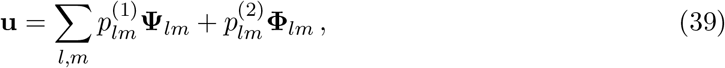

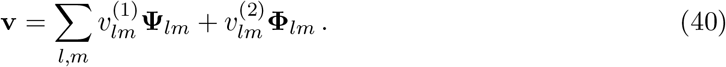

Note that since we are using axis symmetric solutions, all coefficients where *m* ≠ 0 will be equal to zero.

First, from torque balance, we obtain the angular velocity

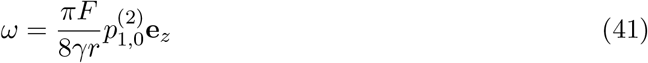

such that the velocity is equal to

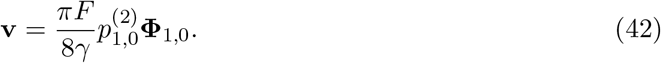

The equations of linear elasticity can be written

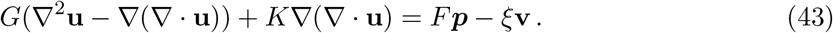

Next, we write our force balance in terms of vector spherical harmonics

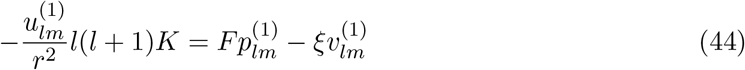

and

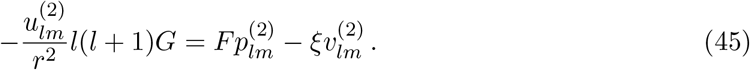

Thus solving for the displacement coefficients we have

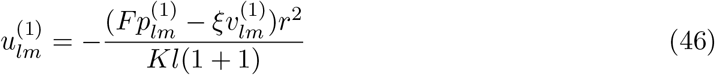

and

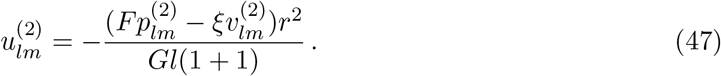

For simplicity, we set the shear modulus *G* = 1. Since cells appear almost incompressible, we set *K* = 100. Finally, we use a traction force magnitude of *F* = 5 which gives a qualitatively similar cellular shear as in experiments. Changing the parameters has little effect on the tilt angle, but does change the magnitude of the shear.

### 5.3 Continuum tilt angle

To measure the resulting tilt angle of cell elongation from the elastic displacement, we determine the deformed shape of initially circular objects on the surface. Each object is initialized as circle with area 1/200th of the area of the sphere to represent the average size of cells in experiments and in vertex model simulations. This defines a set of points **x**_*i*_ that represent a circle. The circle is then deformed according to **X**_*i*_ = **x**_*i*_ + **u**(**x**_*i*_). We then quantify the anisotropy of the deformed circles by fitting ellipses, which defines the tilt angles *β* of elongation.

To obtain a distribution of tilt angles, we place the circle center on the sphere at latitudes between *θ* = 0 and *θ* = *π*, using 1000 points. Additionally, when taking a histogram each angle *θ* contributes a weight 2*πr* sin *θ*, which is the radius of a circle at that latitude. This is because in experimental histoggrams there are more cells near the equator than near the poles which leads to this weight factor.

## 6 Supplementary Figure

**Supplementary Figure 1:**
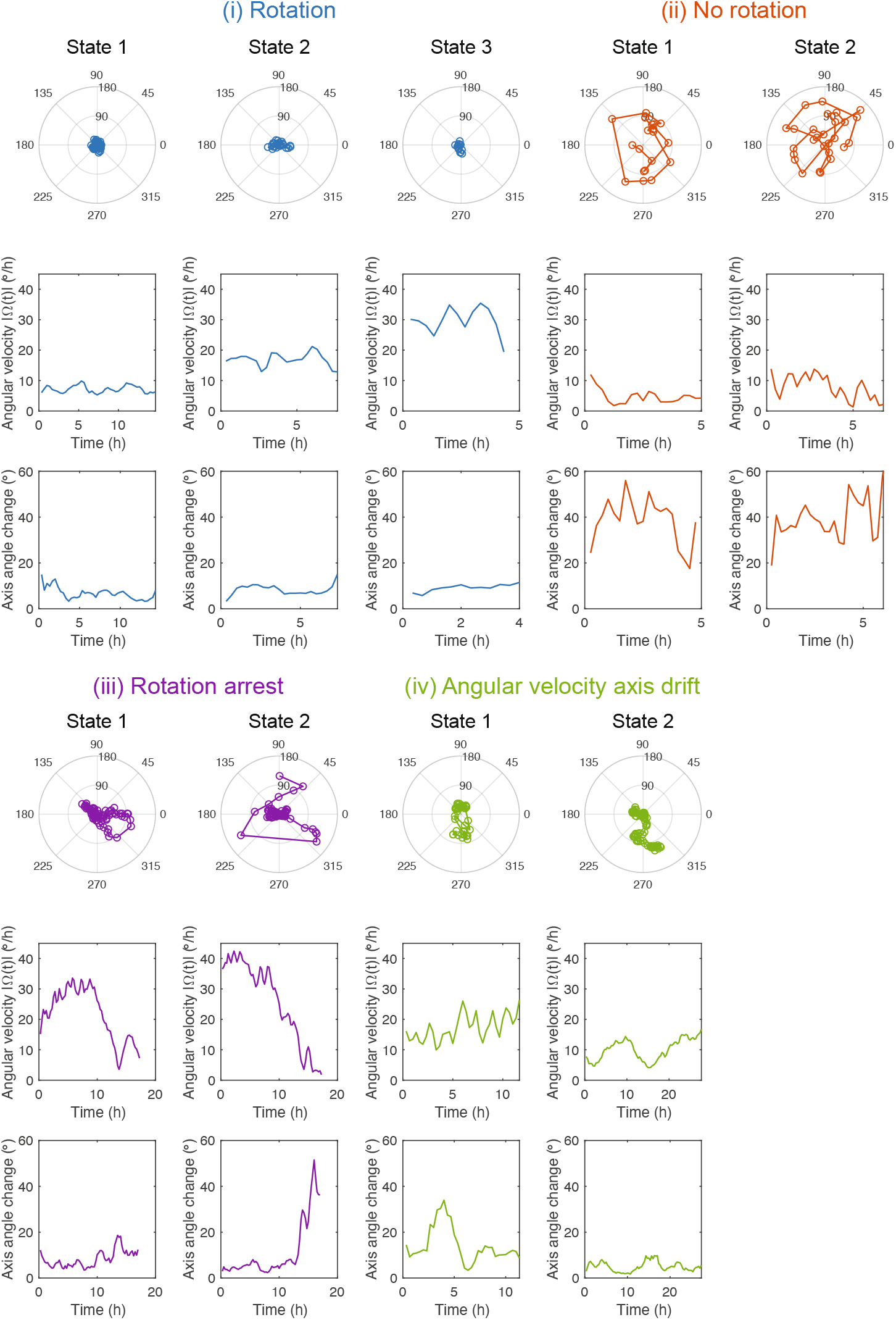
Characterization of rotational dynamics of individual pancreas spheres. There are four distinct types of rotational dynamics observed in experiments: (i) persistent rotation, (ii) no rotation, (iii) rotation arrest, and (iv) rotation axis drift. The first row shows the angular position of the rotation axis in the (*θ, ϕ*) polar plot. The second and third rows show the angular velocity |**Ω**(*t*)| and the axis angle change Δ*ψ/*Δ*t*, respectively.

**Supplementary Figure 2:**
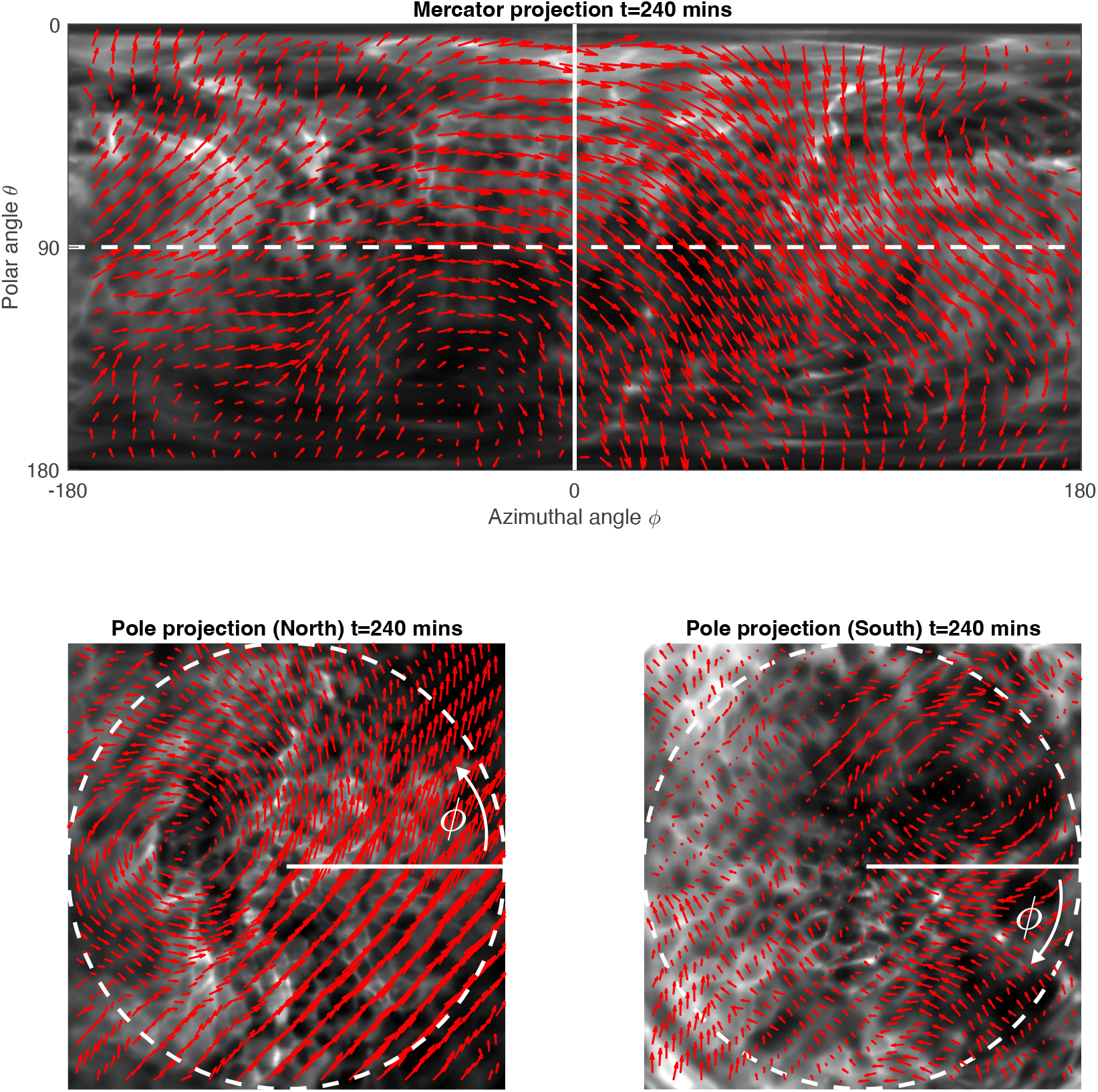
Particle Imaging Velocimetry (PIV) field on the Mercator and pole projections. To obtain the surface velocity field **V**(*θ, ϕ*) at any particular time point *t*, we performed PIV on 3 different projections: (i) Mercator projection (top); (ii) ‘north’ pole projection (bottom left) and (iii) ‘south’ pole projection (bottom right). The PIV fields from the 3 different projections are then stitched together to minimize distortion introduced from different projections. The dotted white line is defined by *θ* = 90° (the ‘equator’). The solid white line is defined by *ϕ* = 0° (the ‘meridian’). See SI Sec. 2.2 for more details.

**Supplementary Figure 3:**
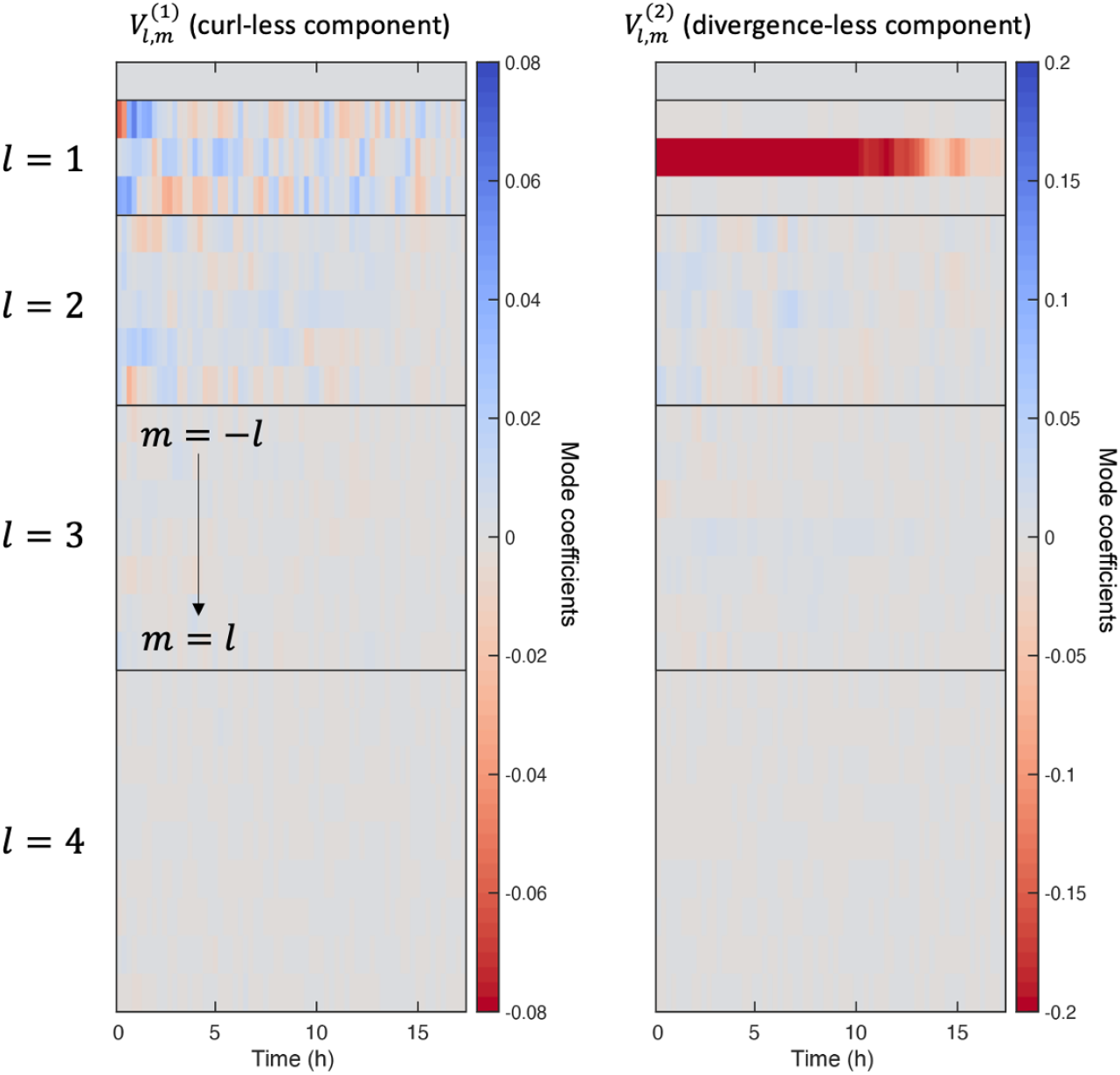
Vectorial spherical harmonics mode coefficients of the velocity field. Visual representation of the mode coefficients 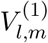 and 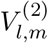 as a function of time, for a sphere undergoing rotation for the first ≈ 10 hours before slowing down. As expected, the amplitude of the 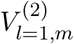 which represents solid body rotation, is the dominant mode.

**Supplementary Figure 4:**
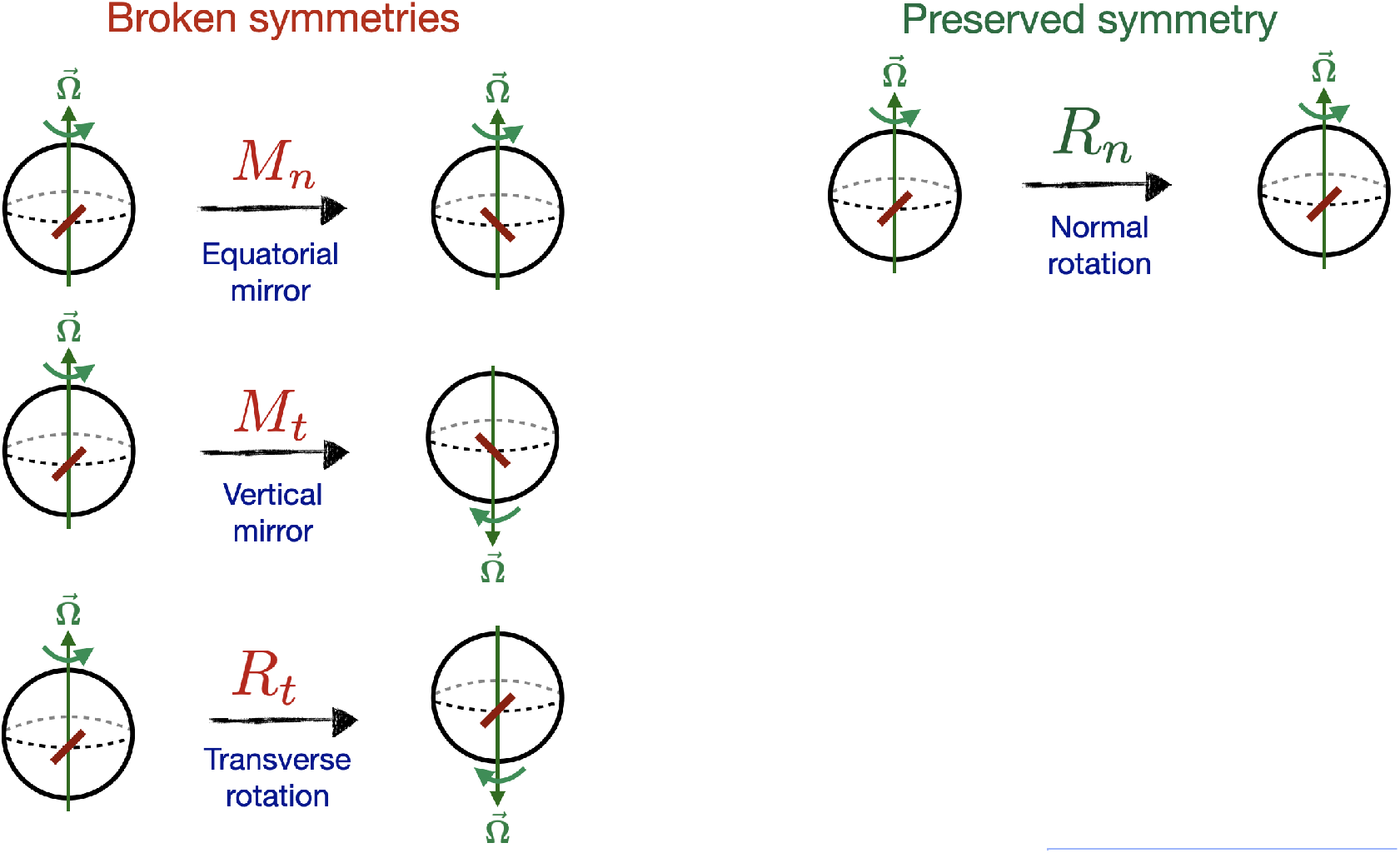
Chiral and up-down symmetry breaking of the cell orientation field on a rotating sphere. Broken (left) and preserved (right) symmetries in a rotating cell sphere with a dominant tilt angle *β* ≈ 45° or *β* ≈ 135° of the cell elongation field.

**Supplementary Figure 5:**
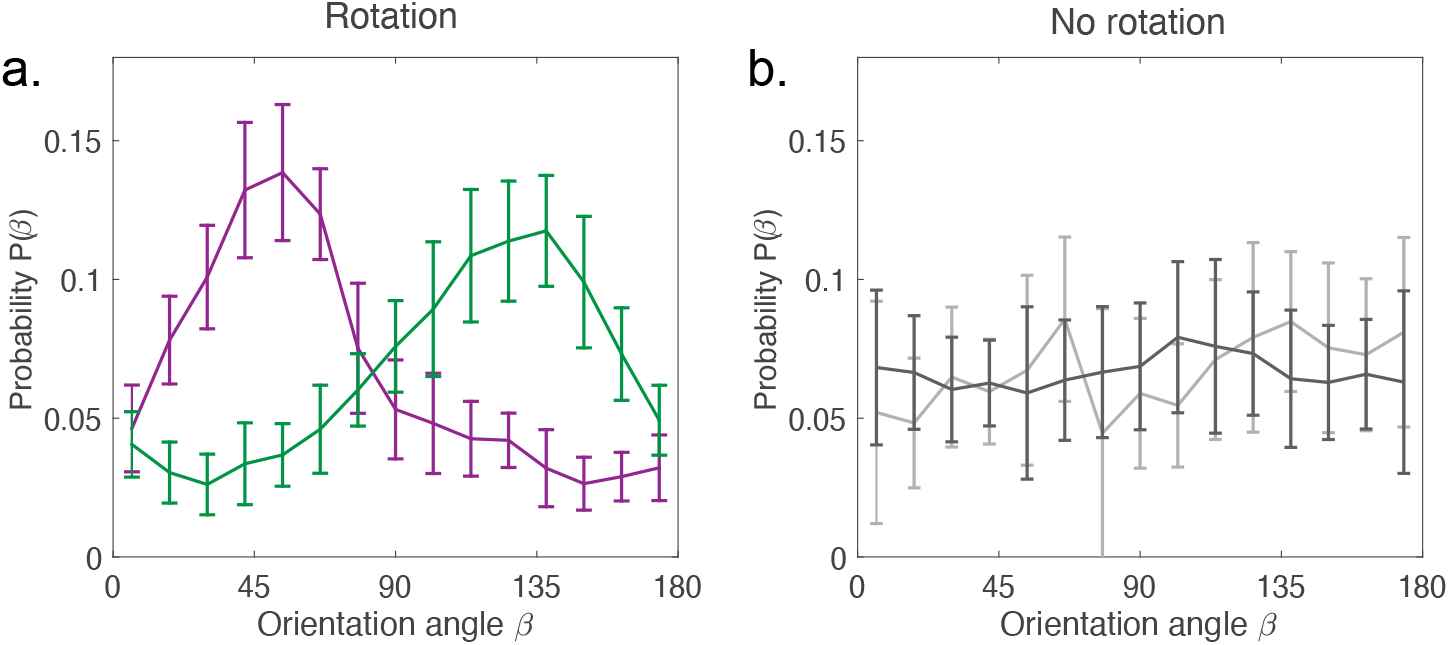
Distribution of cell elongation angles *P* (β) for rotating and non-rotating spheres. The cell shape orientation *β* in rotating pancreas spheres shows a preference to be either ≈ 45° or ≈ 135° for a sustained period of time. This is clearly reflected in the unimodal orientation distribution *P* (*β*) (**a**). In contrast, for non-rotating sphere, the distribution is more or less uniform (**b**).

**Supplementary Figure 6:**
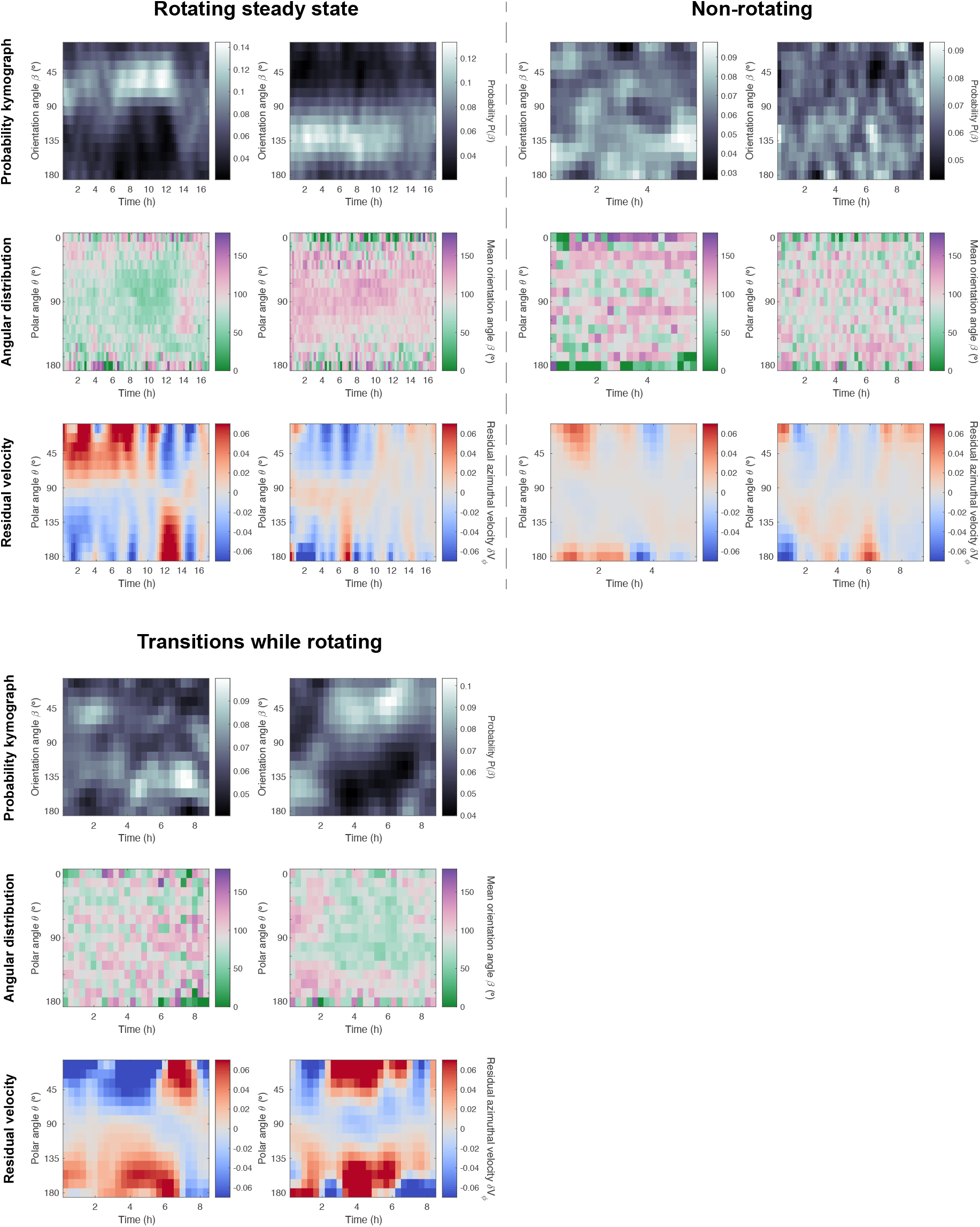
Distribution of cell elongation angle *P* (β) during steady-state rotation (where the dominant tilt angle is constant), transitional rotation (where the dominant tilt angle changes) and for a non-rotating sphere. The first row shows probability-time kymographs of the distribution *P* (*β*). The second row shows azimuthally-averaged cell orientation 〈*β*〉_*ϕ*_(*θ*) as a function of polar angle *θ*. The third row shows the residual azimuthal velocity 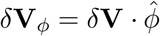 as a function of polar angle *θ*.

**Supplementary Figure 7:**
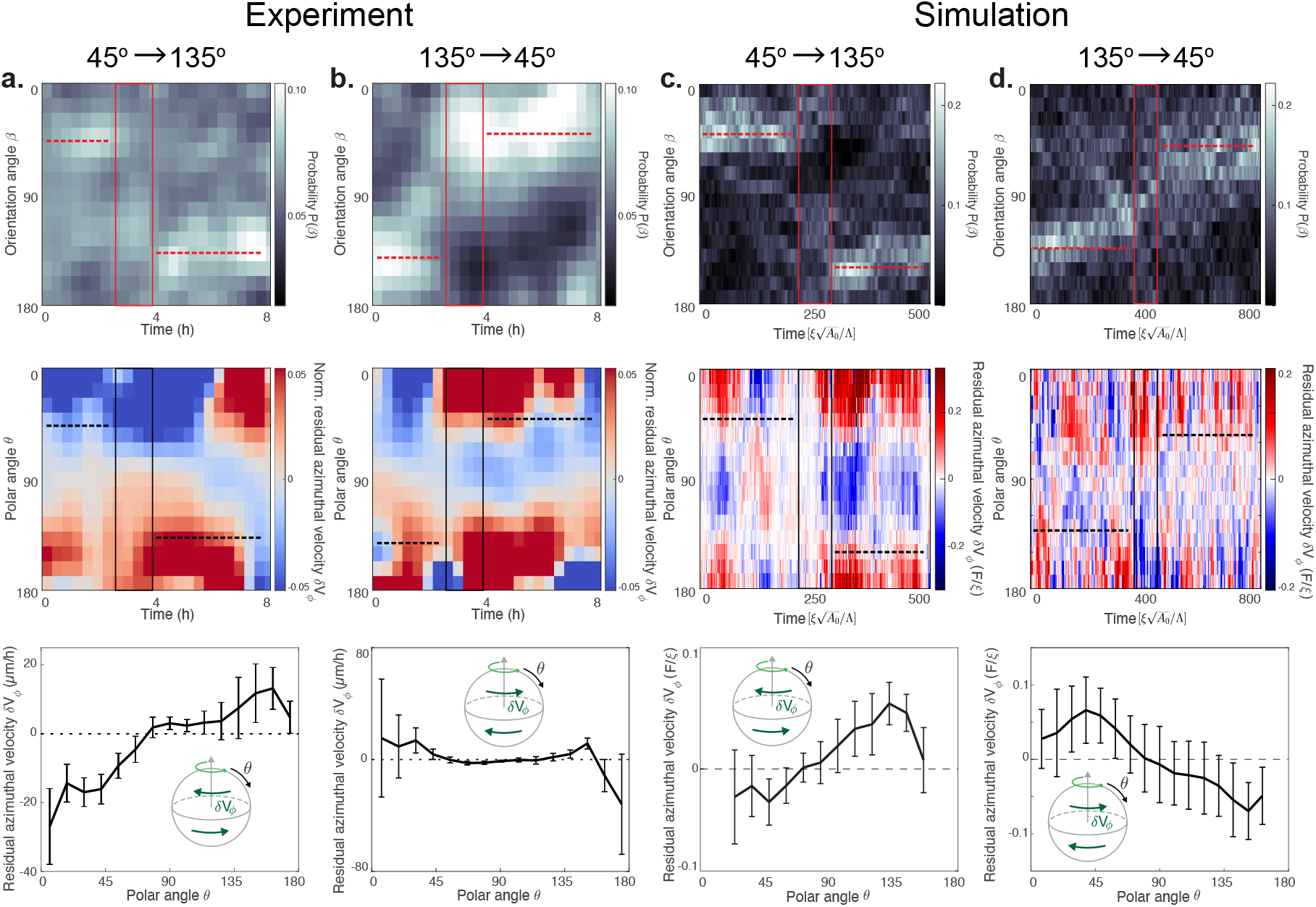
Switch in cell orientation angle β in experiments and simulations. The first row shows the probability-time kymographs of cell orientation angle *β* showing a switch of preferred angle from 45° to 135° (**a,c**) and from 135° to 45° (**b,d**) in both experiments and simulation. The second row shows the probability-time kymographs for the residual azimuthal velocity 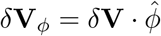. The third row shows the residual azimuthal velocity *δ***V**_*ϕ*_ as a function of polar angle *θ*. Insets show the direction of *δ***V**_*ϕ*_ with respect to rotation axis.

**Supplementary Figure 8:**
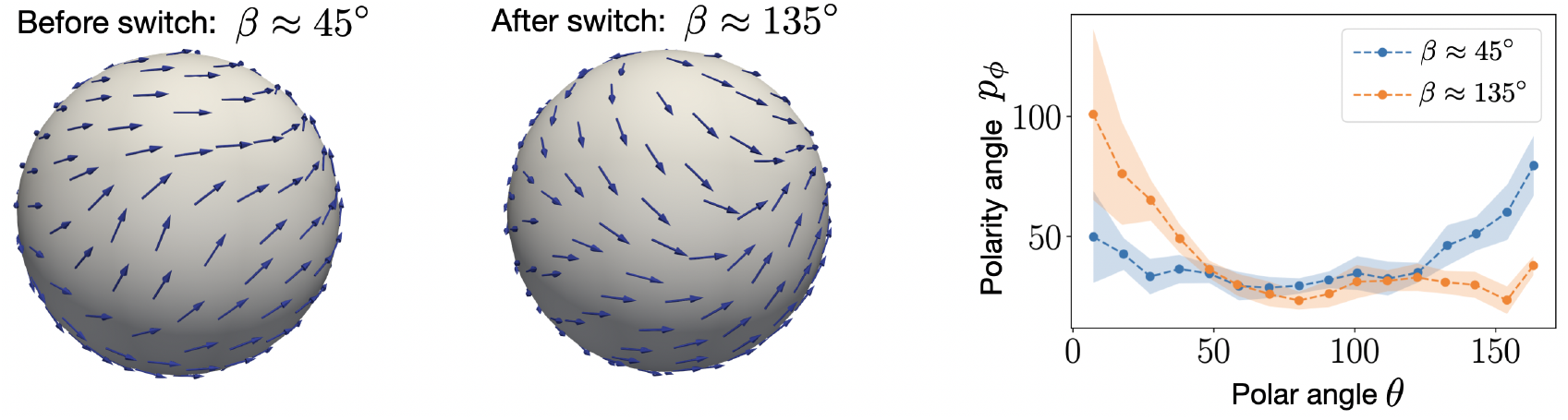
Polarity pattern in active vertex model with switch in the dominant orientation angle. We analyzed the polarity pattern in vertex model simulations with noise in the polarity dynamics. We find that, in contrast to simulations without noise, the peak in the distribution of the cell orientation angle exhibits a switch between *β* ≈ 45° and *β* ≈ 135°. We see a polarity pattern with two topological defects located on the poles. However the nature and positioning of these topological defects seem to change between the time windows with *β* ≈ 45° and *β* ≈ 135°. This is in good agreement with the predictions from our continuum model (see Fig. 5), although we see that the polarity pattern in the vertex model is not entirely axisymmetric. To quantify the polarity pattern, we compute the angle between the polarity field and the azimuthal direction of rotation 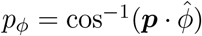 as a function of polar angle *θ*.

**Supplementary Figure 9:**
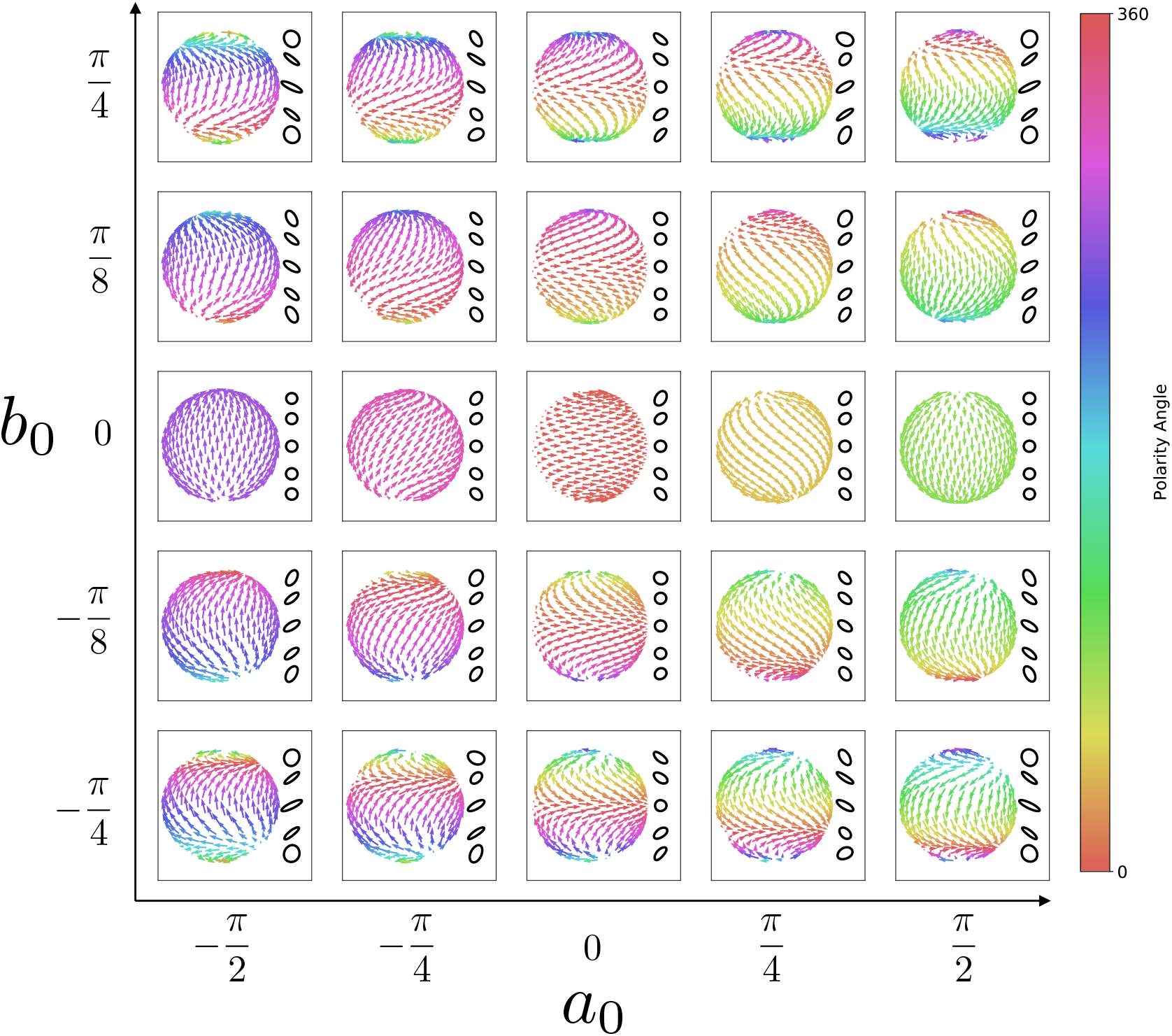
Example steady state solutions from the continuum model. Solutions of the steady state polarity field of the form 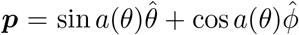 where *a*(*θ*) = *a*_0_ + *b*_0_ ln((1 − cos *θ*)*/*(1 + cos *θ*)), for varying *a*_0_ and *b*_0_. The arrow color indicates the angle *a*(*θ*). Inset: shapes of deformed circles at positions on the sphere. The cell shape orientation *β* only breaks chiral symmetry when *a*_0_ ≠ 0 and *b*_0_ ≠ 0.

## 7 Supplementary Videos

**Video 1 :** Maximum intensity projection of a sphere showing persistent rotation (Fig. 1f(i)). Fluorescence intensity corresponds to membrane dye ROSAmTmG. Time stamp, hh:mm.
**Video 2 :** Maximum intensity projection of a sphere showing no rotation (Fig. 1f(ii)). Fluorescence intensity corresponds to myosin. Time stamp, hh:mm.
**Video 3 :** Maximum intensity projection of a sphere showing rotation arrest (Fig. 1g(i)). Fluorescence intensity corresponds to myosin. Time stamp, hh:mm.
**Video 4 :** Maximum intensity projection of a sphere showing rotation axis drift (Fig. 1g(ii)). Fluorescence intensity corresponds to membrane dye ROSAmTmG. Time stamp, hh:mm.
**Video 5 :** Particle imaging velocimetry (PIV) performed on 3 different projections: (i) Mercator projection (top); (ii) ‘north’ pole projection (bottom left) and (iii) ‘south’ pole projection (bottom right). See SI Sec. 2.2 for the equations defining the mapping. Dotted white line corresponds to *θ* = 90° (i.e. the ‘equator’). The projection is done with respect to lab frame.
**Video 6 :** Simulation of a sphere in the solid rotation regime. A rotating sphere with aligned cell polarities. In this case the correlation length of polarities spans the whole system size.
**Video 7 :** Simulation of a sphere near the yielding transition. Due to an interplay between traction force and the polarity alignment mechanisms, patches of cells with aligned polarities form, but their correlation length is less than the system size. Hence, in this case the rotation axis drifts as patches of aligned polarity cells form and break apart.
**Video 8 :** Simulation of a sphere in the flowing regime. In this case, cell polarity-directed traction forces are uncorrelated. Therefore randomly oriented traction forces lead to continuous flow of cell rearrangements.

## Notes

### Competing Interest Statement

The authors have declared no competing interest.

## References

1. Shellard, A. & Mayor, R. Rules of collective migration: from the wildebeest to the neural crest. Philosophical Transactions of the Royal Society B 375, 20190387 (2020).

2. Muñoz-Dorado, J., Marcos-Torres, F. J., Garcıa-Bravo, E., Moraleda-Muñoz, A. & Pérez, J. Myxobacteria: moving, killing, feeding, and surviving together. Frontiers in microbiology 7, 781 (2016).

3. Copenhagen, K., Alert, R., Wingreen, N. S. & Shaevitz, J. W. Topological defects promote layer formation in Myxococcus xanthus colonies. Nature Physics 17, 211–215 (2021).

4. Stock, J. & Pauli, A. Self-organized cell migration across scales–from single cell movement to tissue formation. Development 148, dev191767 (2021).

5. Brugués, A. et al. Forces driving epithelial wound healing. Nature physics 10, 683–690 (2014).

6. Saw, T. B. et al. Topological defects in epithelia govern cell death and extrusion. Nature 544, 212–216 (2017).

7. Kawaguchi, K., Kageyama, R. & Sano, M. Topological defects control collective dynamics in neural progenitor cell cultures. Nature 545, 327–331 (2017).

8. Livshits, A., Shani-Zerbib, L., Maroudas-Sacks, Y., Braun, E. & Keren, K. Structural inheritance of the actin cytoskeletal organization determines the body axis in regenerating hydra. Cell reports 18, 1410–1421 (2017).

9. Hoffmann, L. A., Carenza, L. N., Eckert, J. & Giomi, L. Theory of defect-mediated mor-phogenesis. Science advances 8, eabk2712 (2022).

10. Mongera, A. et al. A fluid-to-solid jamming transition underlies vertebrate body axis elon-gation. Nature 561, 401–405 (2018).

11. Saadaoui, M., Rocancourt, D., Roussel, J., Corson, F. & Gros, J. A tensile ring drives tissue flows to shape the gastrulating amniote embryo. Science 367, 453–458 (2020).

12. Keber, F. C. et al. Topology and dynamics of active nematic vesicles. Science 345, 1135–1139 (2014).

13. Ellis, P. W. et al. Curvature-induced defect unbinding and dynamics in active nematic toroids. Nature Physics 14, 85–90 (2018).

14. Shankar, S., Bowick, M. J. & Marchetti, M. C. Topological sound and flocking on curved surfaces. Physical Review X 7, 031039 (2017).

15. Hirata, E., Ichikawa, T., Horike, S.-i. & Kiyokawa, E. Active K-RAS induces the coherent rotation of epithelial cells: a model for collective cell invasion in vitro. Cancer science 109, 4045–4055 (2018).

16. Chin, A. S. et al. Epithelial cell chirality revealed by three-dimensional spontaneous rotation. Proceedings of the National Academy of Sciences 115, 12188–12193 (2018).

17. Brandstätter, T. et al. Curvature induces active velocity waves in rotating multicellular spheroids. arXiv preprint arXiv:2110.14614 (2021).

18. Barlan, K., Cetera, M. & Horne-Badovinac, S. Fat2 and Lar define a basally localized planar signaling system controlling collective cell migration. Developmental Cell 40, 467–477 (2017).

19. Tanner, K., Mori, H., Mroue, R., Bruni-Cardoso, A. & Bissell, M. J. Coherent angular motion in the establishment of multicellular architecture of glandular tissues. Proceedings of the National Academy of Sciences 109, 1973–1978 (2012).

20. Wang, H., Lacoche, S., Huang, L., Xue, B. & Muthuswamy, S. K. Rotational motion during three-dimensional morphogenesis of mammary epithelial acini relates to laminin matrix assembly. Proceedings of the National Academy of Sciences 110, 163–168 (2013).

21. Fernández, P. A. et al. Surface-tension-induced budding drives alveologenesis in human mammary gland organoids. Nature Physics 17, 1130–1136 (2021).

22. Hof, L. et al. Long-term live imaging and multiscale analysis identify heterogeneity and core principles of epithelial organoid morphogenesis. BMC biology 19, 1–22 (2021).

23. Greggio, C. et al. Artificial three-dimensional niches deconstruct pancreas development in vitro. Development 140, 4452–4462 (2013).

24. Farhadifar, R., Röper, J.-C., Aigouy, B., Eaton, S. & Jülicher, F. The influence of cell mechanics, cell-cell interactions, and proliferation on epithelial packing. Current Biology 17, 2095–2104 (2007).

25. Bi, D., Yang, X., Marchetti, M. C. & Manning, M. L. Motility-driven glass and jamming transitions in biological tissues. Physical Review X 6, 021011 (2016).

26. Popović, M., Druelle, V., Dye, N. A., Jülicher, F. & Wyart, M. Inferring the flow properties of epithelial tissues from their geometry. New Journal of Physics 23, 033004 (2021).

27. Honda, H., Yamanaka, H. & Dan-Sohkawa, M. A computer simulation of geometrical configurations during cell division. Journal of theoretical biology 106, 423–435 (1984).

28. Dye, N. A. et al. Self-organized patterning of cell morphology via mechanosensitive feedback. Elife 10, e57964 (2021).

29. Yang, Q. et al. Cell fate coordinates mechano-osmotic forces in intestinal crypt formation. Nature Cell Biology 23, 733–744 (2021).

30. Gehrels, E. W., Chakrabortty, B., Merkel, M. & Lecuit, T. Genetic and geometric heredity interact to drive polarized flow in the Drosophila embryo. bioRxiv (2022).

31. Shinohara, K. & Hamada, H. Cilia in left–right symmetry breaking. Cold Spring Harbor perspectives in biology 9, a028282 (2017).

32. Taniguchi, K. et al. Chirality in planar cell shape contributes to left-right asymmetric epithelial morphogenesis. Science 333, 339–341 (2011).

33. Hadidjojo, J. & Lubensky, D. K. Spontaneous Chiral Symmetry Breaking in Planar Polarized Epithelia. arXiv preprint arXiv:1708.08560 (2017).

## References

1. Muzumdar, M. D., Tasic, B., Miyamichi, K., Li, L. & Luo, L. A global double-fluorescent Cre reporter mouse. genesis 45, 593–605 (2007).

2. Greggio, C. et al. Artificial three-dimensional niches deconstruct pancreas development in vitro. Development 140, 4452–4462 (2013).

3. Berg, S. et al. Ilastik: interactive machine learning for (bio) image analysis. Nature methods 16, 1226–1232 (2019).

4. Thielicke, W. & Sonntag, R. Particle Image Velocimetry for MATLAB: Accuracy and enhanced algorithms in PIVlab. Journal of Open Research Software 9(2021).

5. Aigouy, B., Cortes, C., Liu, S. & Prud’Homme, B. EPySeg: a coding-free solution for automated segmentation of epithelia using deep learning. Development 147, dev194589 (2020).

6. Farhadifar, R., Röper, J.-C., Aigouy, B., Eaton, S. & Jülicher, F. The influence of cell mechanics, cell-cell interactions, and proliferation on epithelial packing. Current Biology 17, 2095–2104 (2007).

7. Honda, H., Yamanaka, H. & Dan-Sohkawa, M. A computer simulation of geometrical configurations during cell division. Journal of theoretical biology 106, 423–435 (1984).

